# RNabel—A Standalone Software Tool for Annotating Tandem Mass Spectra of Modified Ribonucleic Acids

**DOI:** 10.64898/2026.06.22.733900

**Authors:** Ge Song, Yi-Jie Nathan Du, Rui-Xiang Sun, Meng-Qiu Dong

**Affiliations:** Graduate School of Peking Union Medical College & Chinese Academy of Medical Sciences, Beijing 100730, China; National Institute of Biological Sciences, Beijing, Beijing 102206, China; Tsinghua Institute of Multidisciplinary Biomedical Research, Tsinghua University, Beijing 100084, China; Bachelor of Science in Computer Science program, University of London

**Keywords:** RNA and ssDNA modifications, tandem mass spectrometry, MS/MS spectrum annotation, nucleic acids, software tool, automated analysis, RNabel

## Abstract

Ribonucleic acid (RNA) modifications, with over 170 identified types, play diverse roles in cellular processes. The past decade has witnessed surging demand for accurate identification and localization of RNA modifications in both endogenous and synthetic therapeutic RNAs. With accurate spectral annotation for RNA, tandem mass spectrometry (MS/MS) can meet this demand. Here we present RNabel, a user-friendly software tool for in-depth annotation of MS/MS spectra of RNA oligonucleotides. RNabel considers a full set of backbone-cleavage ions (*a*, *b*, *c*, *d*, *a-B*, *w*, *x*, *y*, *z*) in which the ribonucleotide unit could be A, U, C, G, Y (pseudouridine), or I (Inosine). Additionally, RNabel considers 196 modifications on the base, the phosphoribose linkage, the 5′ or the 3′ terminus, or detachment of a sub-nucleotide fragment as a neutral or charged group. Users can create new components if needed, including ribonucleotides, modifications, neutral or charged groups that could detach from a ribonucleotide. RNabel efficiently processes large datasets in four acceptable formats including .mgf, .raw, .txt from msConvert, and RNabel batch files. Multiple statistical metrics are provided for quality assessment of spectral annotation. To accelerate RNA modification analysis, RNabel is made freely available for Mac and Windows users at https://github.com/songge1111/RNabel/releases.

**Graphic Abstract:** 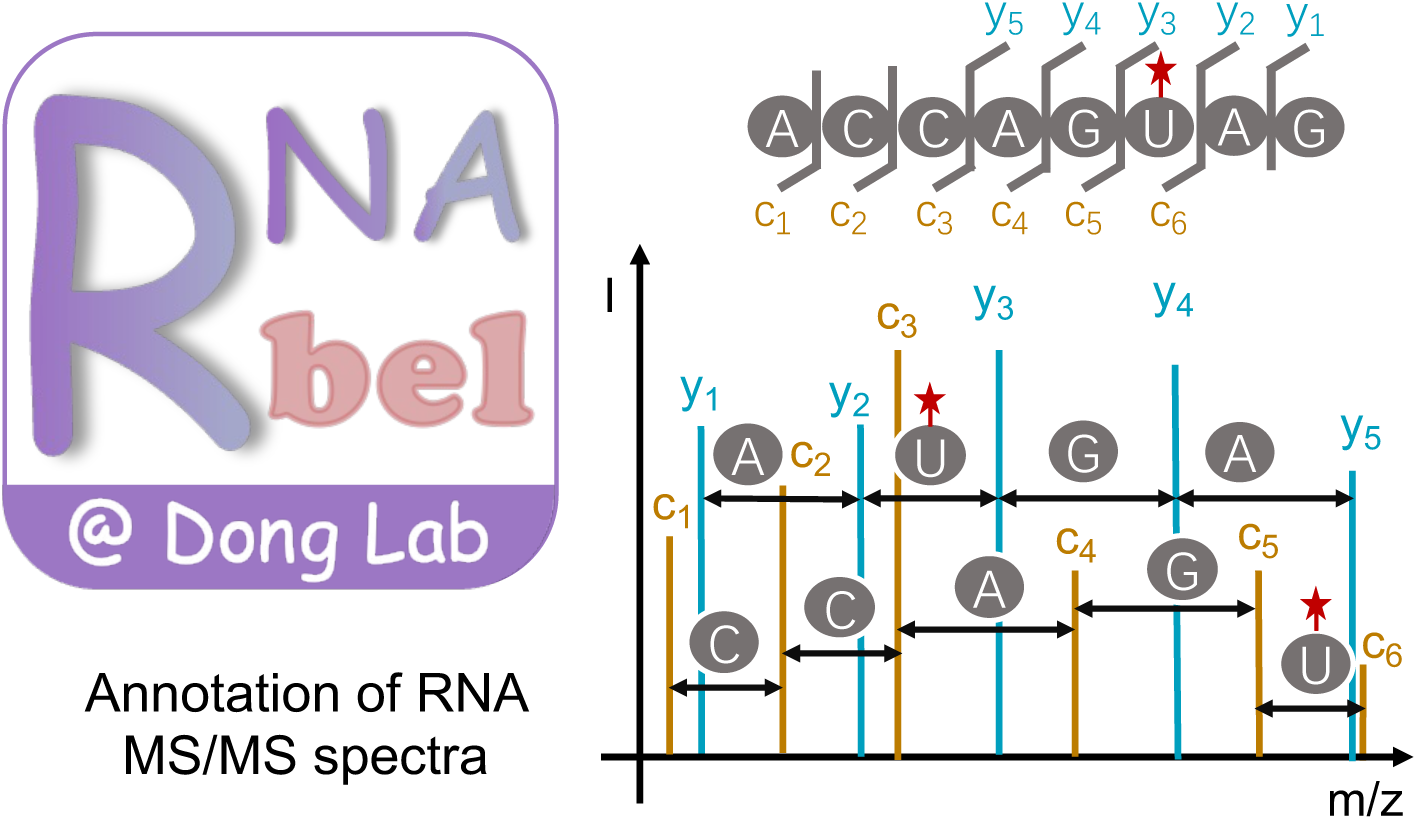

## Introduction

RNA modifications have been shown to influence the 3D structure, stability, subcellular localization, and function of RNA molecules across diverse biological contexts^1^. To date, over 170 distinct types of RNA modifications have been identified in prokaryotic and eukaryotic organisms^2,3^. These modifications constitute a new layer of gene-expression regulation, termed epitranscriptome^1^, with demonstrated roles in key cellular processes such as translation, splicing, and immune recognition^1,4,5^. Incorrect deposition of RNA modification may lead to abnormal development^6^ and cancer^7,8^. For example, METTL1-mediated m^7^G (7-methylguanosine) is required for the production of mature *let-7e* miRNA, which blocks cancer metastasis by suppressing *HMGA2* (high mobility group AT-hook2)^9^.

In addition, RNA modifications are indispensable for RNA-based therapeutics, such as mRNA vaccines, which have helped the world fight against the SARS-CoV2 pandemic and for which a Nobel prize was awarded in 2023^10,11,12^. To enhance the stability to mRNA vaccine in the human body and to minimize unwanted immunogenicity towards the vaccine mRNA, pseudouridine modifications are introduced routinely by chemical synthesis.

Several experimental approaches have been developed to detect RNA modifications. Next-generation sequencing-based methods, such as MeRIP-Seq^13^ and BACS^14^, infer modification sites by exploiting reverse transcription signatures and antibody enrichment. While suitable for transcriptome-scale profiling, these techniques suffer from limited resolution, antibody dependence, and ambiguity regarding the nature of modification. Additionally, these methods detect only one RNA modification type per experiment.

Recently, nanopore-based direct RNA sequencing has emerged as a single-molecule strategy that identifies modifications via ionic current deviations^15^. However, this signal is additionally influenced by flanking nucleotides (k-mer context)^15,16^. Therefore, extensive training is required for each modification to be analyzed^16,17^. Consequently, while being a promising method to look forward to, nanopore-based sequencing is faced with many challenges.

In contrast, mass spectrometry (MS) can detect different RNA modifications in one experiment. To data, this is carried out by digesting RNA molecules into mononucleotides or mononucleosides prior to MS analysis. Thus, although it is a sensitive and quantitative assay, the sequence context of detected modifications is lost^18,19,20^.

Tandem mass spectrometry (MS/MS) has emerged as a powerful method for sequence and modification analysis of RNA molecules (see Figure 1). In this method, RNA molecules are fragmented in the gas phase along the phosphoribose backbone, resulting in sequencing ions. The fragmentation pattern of oligonucleotides by CID was first reported in 1992^21,22,23,24,25^. This pioneering study defined characteristic ion series including *a, b, c, d-ions* from the 5′ end, *w, x, y, z*-*ions* from the 3′ end, and *a-B* ions that correspond to *a-ions* losing a base (see Figure 1B). From the charge and mass-to-charge ratios (m/z) of these fragment ions, the full sequence of the parent RNA ion may be deduced. If an RNA is chemically modified, a characteristic mass shift is expected for fragment ions carrying the modification(s). Based on such information, the modification group(s) can be identified and located (see Figure 1C).

**Figure 1:**
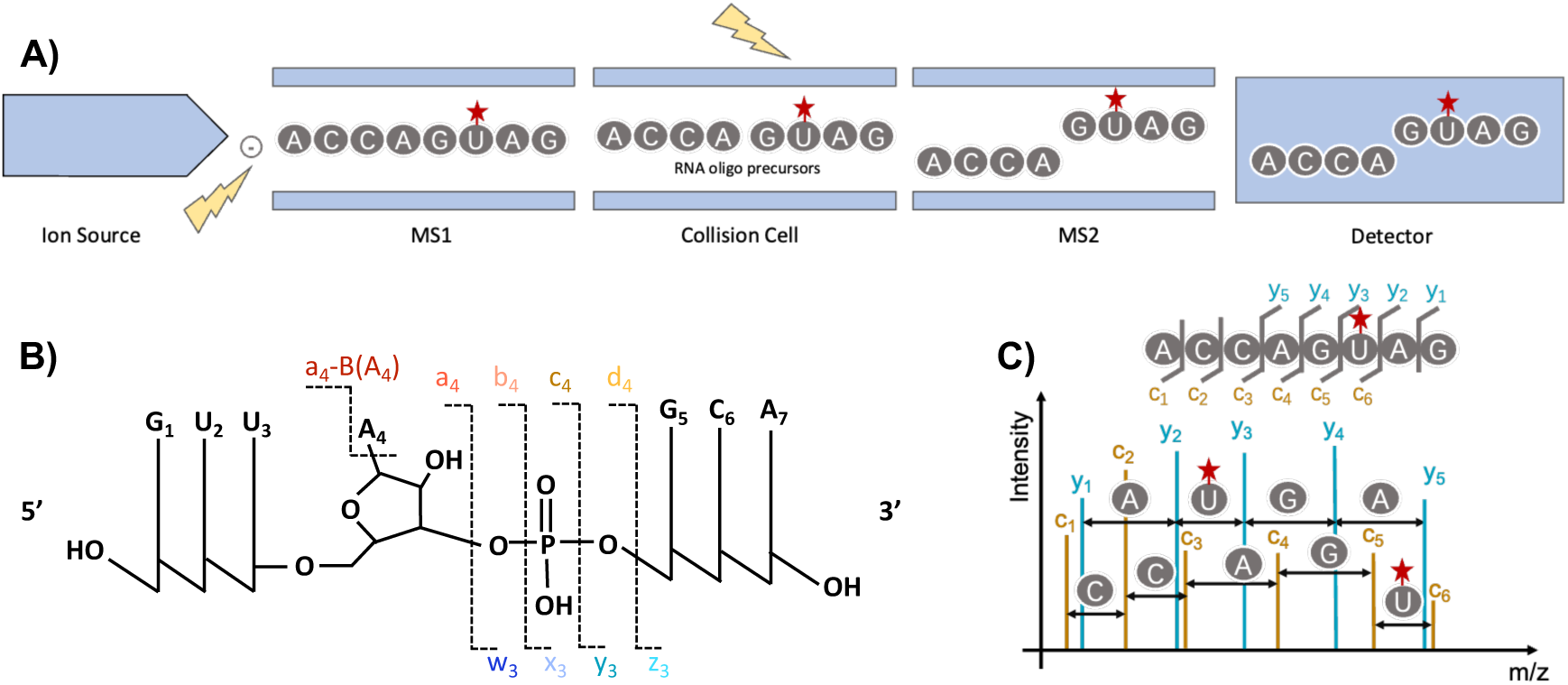
Identification of RNA sequence and modification based on MS/MS analysis. A) Illustration of MS/MS analysis of RNA oligonucleotides. B) RNA fragmentation sites and nomenclature of corresponding ion types proposed by McLuckey et al^23^. The 5′-fragments include *a*, *b*, *c*, and *d* ions, and the 3’-fragments include *w*, *x*, *y*, and *z*-ions. If an *a*-ion has a base detached from a ribose, it becomes an a-B ion. C) Localization of an RNA modification through a modification-specific mass shift in sequencing ions.

MS/MS analysis has been utilized to characterize various types of RNAs, including the endogenous RNAs (e.g., tRNA) and the synthetic ones (e.g., mRNA vaccines)^26,27^. However, the need for flexible and comprehensive annotation of RNA MS/MS spectra remains unaddressed, especially for RNAs carrying unknown or synthetic, unconventional modifications. In published MS/MS spectra of modified RNAs, many peaks are not assigned (one example is shown in Figure S1). Close examination of the unassigned peaks revealed to us that many of them corresponded to sequence-informative fragment ions. Although experts can manually assign identity to some of them, it is impractical for high-throughput MS/MS data, and poses a formidable barrier for newcomers to the field.

Among the available software tools for annotating RNA MS/MS spectra, some are not freely available (e.g., Biopharma Finder, Byonic), some lack flexibility as they are imbedded in a search engine (e.g., Ariadne^28^, RNAModMapper^29^, NASE^26^, Pythease^27^) and require predefined modifications and known sequences. A handful of interactive freewares (e.g., OMA/OPA^30^, RoboOligo^31^, Aom2S^32^, FAST MS^33^) offer limited support for user-defined modifications and batch processing. To address these limitations, we present RNabel, a highly configurable software platform for annotating RNA MS/MS spectra with high peak interpretation rates. RNabel allows users to define custom RNA modifications, nucleotide residues, 5′ and 3′ terminal groups, and sub-nucleotide fragments, thus facilitating exploration of unassigned spectral peaks. This function is also needed for understanding novel modifications and complex fragmentation patterns. RNabel operates independently of any specific instrument platform, enabling seamless handling of individual MS/MS spectra as well as large-scale LC-MS/MS datasets through automated batch processing. With flexible customization and quantitative annotation metrics, RNabel is designed to support both routine analysis and exploratory studies. We expect RNabel to accelerate mass spectrometry method development for RNA analysis.

## Materials and Methods

### Materials

To develop and evaluate the RNabel annotation pipeline, we used two in-house MS/MS datasets. Dataset #1, used for spectral presentation, contains MS2 spectra of RNA oligonucleotides carrying different modifications (e.g., m⁶A, m^6^Am) and terminal groups. Dataset #2, used for evaluation, contains 333 MS/MS spectra generated by 19 unmodified RNA oligonucleotides. The oligonucleotides (Table S3) were synthesized and HPLC-purified by Accurate Biotechnology (Hunan) Co., Ltd..

### Software Implementation and Architecture

Implemented in Python 3.9 with a PySide6-based GUI, RNabel is designed as a standalone, cross-platform tool that is compatible with both macOS and Windows operating systems. RNabel is built on an architecture of modular design, which affords upgrade flexibility and ease of maintenance. The core components of the RNabel software are as follows.

**1) Spectrum pre-processing module**: This module performs initial data processing, including noise filtering and isotope detection, to ensure that the input mass spectrometry data is suitable for further analysis.
**2) Theoretical fragment ion calculator module**: This component generates theoretical fragment ions with information specified by users, which includes RNA/DNA sequence, modifications, 5′/3′ termini, and the theoretical ion types to be considered.
**3) Matching module**: The matching module matches experimental peaks with theoretical fragment ions within a user-defined mass tolerance (ppm) window. Additional parameters, such as fragmentation method (CID/HCD or ETD) and intensity thresholds, are used to refine the matching process. This ensures accurate identification and annotation of peaks within the spectra.
**4) Statistics module**: The statistics module in RNabel provides a set of statistical metrics to inform users about the quality of spectral annotation. These metrics include portion of summed intensities of interpreted peaks, the portion of interpreted peak counts, and sequence coverage, etc. (Table S1) The formulas for calculating these metrics are detailed in Table S1.
**5) Graphical annotation renderer**: After successful peak matching, this module generates high-resolution, color-coded graphical representations of the annotated spectra, leveraging Matplotlib and Seaborn for visualization. These visualizations include fragmentation maps, annotated peak identities, and mass errors.

RNabel is distributed under a proprietary license and is freely available for academic research purposes. Modification and redistribution of the source code are prohibited. For commercial use or integration into proprietary software, please contact the authors to discuss licensing arrangements.

### Fragment Ion Calculation and Annotation Strategy

RNabel supports comprehensive calculation of theoretical fragment ions, including *a-B, a, b, c, d, w, x, y, z, M* (parental ions), *s-B* (including all sequencing ions that have a loss of a base or other chemical groups, except *a-B*), *i* (internal ions), and *B* (low m/z range peaks) ions (see Table 1 for ion codes and Figure 3 for their classification). When needed, modified residues are taken into consideration with position-specific mass shifts. RNabel supports all published RNA modifications listed on the MODOMICS website (https://genesilico.pl/modomics/) and allows users to define new modifications.

**Table 1.** Ion types supported by RNabel and their codes. RNabel assigns codes to five categories of fragment ions used for RNA/DNA tandem MS spectrum annotation. Sequencing ions (*a*, *b*, *c*, *d*, *a-B*, *w*, *x*, *y*, *z*) are collectively represented by the code “s” in the visualization module, while each subtype retains its standard nomenclature based on the convention established by McLuckey et al.^23^. Other ion types, including low m/z range peaks (B), parent peaks (M), internal ions (i), and sequencing ions with base loss (s-B), are also assigned single-letter codes as listed.

| Ion types in RNA tandem MS spectra | Code | Note |
| --- | --- | --- |
| Sequencing ions | s | Reported individually as *a, b, c, d, w, x, y, z*, and *a-B* |
| Low m/z range peaks | B |  |
| Parent peaks | M |  |
| Internal ions | i |  |
| Sequencing ions with base loss | s-B |  |

The annotation engine matches observed peaks to predicted ions within a user-defined mass tolerance window (default: +− 20 ppm). As illustrated in Figure S2, some ions share the same m/z values. To minimize ambiguity, RNabel considers fore priority levels as follows:

**1) First priority**: *a-B, a, w, c, y, M, B* ion.
**2) Second priority**: *b, d, x, z* ions.
**3) Third priority**: *i* ions.
**4) Forth priority**: *s-B* ions.

This is based on the fragmentation characteristics of RNA oligos^34^. By default, RNabel includes first-priority ions, and users can enable the second-, third-, and forth-priority ions. Additionally, ions within the same priority group can be selected separately for single ion-type matching.

To ensure publication-quality output, RNabel generates the following results:

1. Spectrum plots with labeled ions.
2. Error plots for visualizing the mass difference of observed and theoretical ions in ppm.
3. Fragmentation maps representing the backbone coverage of RNA/DNA oligos.
4. Optionally, tables of matched peaks in mgf format including statistical parameters, can be downloaded.
5. A list of m/z values of theoretical fragments.

To further validate the results, RNabel provides the option for rechecking annotations.

### Memory Management and Multi-File Loading

RNabel employs a memory management strategy designed to efficiently load multiple MS data files without exceeding available memory resources. This is achieved through a multi-level indexing strategy, enabling simultaneous reading of multiple data files while optimizing memory usage.

The process begins by sequentially reading the raw files from a list. Each file is divided into blocks of a maximum of 1000 spectra, which are stored as temporary files. Each spectrum is indexed with several key attributes: {File ID: {Spectrum ID: {block ID, start, length}}}, where the File ID refers to the raw file ID, and Spectrum ID is the scan number. This indexed information is stored in a separate index file, enabling RNabel to retrieve any spectrum with O(1) time complexity when needed.

This multi-level indexing strategy, coupled with binary storage of spectra, allows RNabel to handle large datasets efficiently, preventing memory overload while providing rapid access to any spectrum via direct indexing. See Figure S3 for a diagram illustrating this strategy.

### Batch Spectrum Annotation in RNabel

To use the batch annotation function, users need import a CSV file containing essential spectrum information and identity details. The required column headers are summarized in Table S2. This batch file can be generated using the RNAsogo (a search engine for RNA MS/MS data to be published separately). Once the file is loaded, RNabel automatically displays the annotated spectrum as users switch between spectra, thereby enhancing the efficiency of large-scale spectrum analysis. For more details on the batch annotation workflow, see Figure S4.

### Isotopic Distribution Simulation

To better understand the mass spectra of RNA, we simulated the isotopic distribution patterns of 4–116 nt RNA oligos, considering all possible nucleotide compositions. For each composition, the theoretical isotopic distribution was calculated using the EMASS algorithm^35,36^. The most intense peak within each cluster was identified and its relationship with oligonucleotide length or mass is shown in Figure 8A.

### Unsupervised Threshold Determination via Gaussian Mixture Modeling

To objectively establish a cosine-similarity threshold separating genuine RNA ions from random noise, an unsupervised Gaussian Mixture Model (GMM) was applied to all candidate assignments from the full-library search of Dataset #2. A total of 248,564 candidates falling within a mass tolerance window of ±20 ppm are represented as two-dimensional feature vectors (mass error in ppm, isotope-distribution cosine similarity) and standardized to zero mean and unit variance prior to fitting (Figure S13). A three-component GMM with full covariance matrices was selected based on a local minimum in the Bayesian Information Criterion (BIC) at k = 3 and the physical interpretability of the resulting clusters (True Match, Gray Zone, and Noise). Model parameters were estimated by Expectation-Maximization (EM) with five K-Means restarts (n_init = 5, max_iter = 300; scikit-learn), and robustness was confirmed by repeating the fit across 20 independent random seeds, all of which converged to essentially identical component parameters. The Bayesian posterior probability P(True Match | x) exhibited a steep sigmoidal transition near cosine = 0.8, coinciding with a valley in the cosine-similarity distribution, and this value was therefore adopted as the filtering threshold.

## Results

### RNabel Workflow Overview

RNabel is a standalone software tool for annotating MS/MS spectra of RNA or single-stranded DNA by matching experimental peaks with theoretical fragment peaks. With user-provided input files of MS/MS spectra and information about RNA sequence, modification, ion mode, etc. RNabel performs annotation in four steps (see Figure 2):

1. **Spectrum File Reading and Peak Extraction**: RNabel reads the input MS data files and extracts peak lists for each MS/MS spectrum.
2. **Theoretical Fragment Peaks Calculation**: Theoretical fragment peaks are calculated based on the input RNA sequence and modification(s).
3. **Peak Matching**: Experimental peaks are matched to theoretical peaks based on the default or user-defined intensity threshold within a user-defined mass tolerance window.
4. **Annotation and Visualization**: Matched peaks are colored to represent different ion types. Mass errors and the fragmentation map are also visualized, with all outputs available for download in vector graphic format.

**Figure 2:**
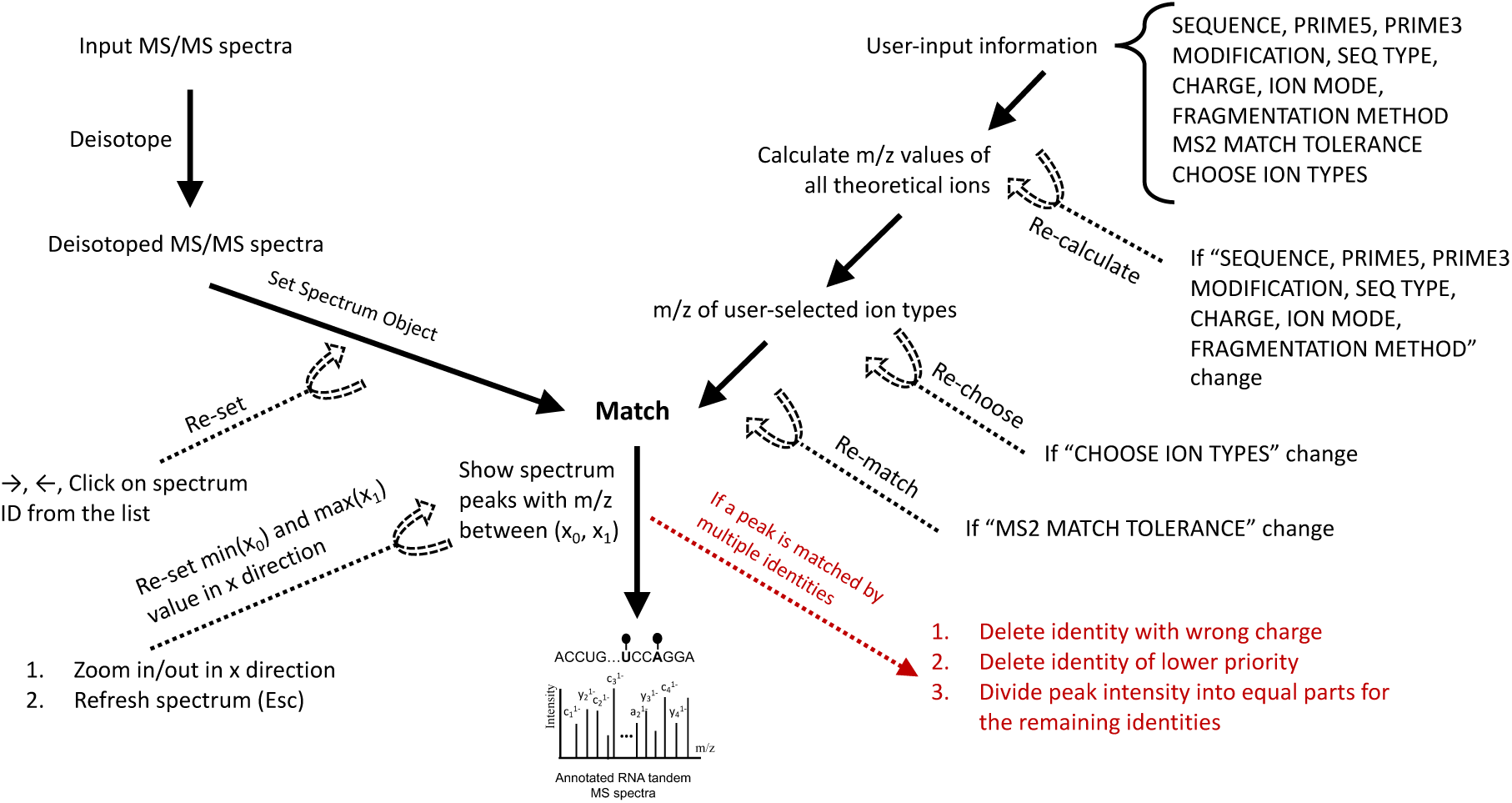
Computational workflow of RNabel. RNabel annotation starts with input MS/MS files (upper left) and the necessary user-input information (upper right) about RNA sequence, 5′- and 3′-terminal groups, modifications, etc…. Mass tolerance, fragmentation method, and ion type selection have default values but can be customized if desired. RNabel constructs a spectrum object and can deisotope, if this option is activated, to simplify the peak list before calculating theoretical fragment ions and matching them to experimental *m/z* peaks within the specified tolerance. RNabel also supports interactive zoom-in and zoom-out operations to facilitate detailed inspection of specific spectral regions. When a peak is matched to multiple candidate identities, RNabel applies a conflict-resolution mechanism by removing identities with incorrect charge, excluding lower-priority identities, and distributing peak intensity equally among the remaining valid identities. Whenever the sequence information, selected ion types, or mass tolerance is modified, RNabel automatically recalculates or re-filters the theoretical fragments and re-performs peak matching accordingly. The final outputs include annotated spectra, fragmentation maps, error plots, and statistical metrics such as sequence and position coverage, all of which can be exported in publication-quality formats.

In addition to the standard workflow, RNabel supports batch annotation by reading the sequence information of each spectrum from a RNabel batch file, improving efficiency for large-scale spectrum analysis. For detailed information on batch spectrum annotation, see the Methods section.

### Ion Classification

RNabel classifies theoretical fragment peaks into five categories, each assigned a specific code (*s, B, M, i, s-B*) for visualization (see Table 1). These categories include:

1. **Sequencing Ions (*s*)**: fragment ions produced from a single backbone cleavage event, providing sequence information crucial for sequence identification (Figure 3A-B).
2. **Low m/z Range Peaks (*B*)**: ions appearing in the lower m/z range of a MS/MS spectrum, offering additional fragment information useful for spectrum interpretation (Figure 3C).
3. **Internal Ions (*i*)**: ions formed through two or more backbone cleavages (Figure 3D).
4. **Parent Peaks (*M*)**: intact precursor ions and precursor ions that have lost one or more small chemical groups while retaining an intact backbone (Figure 3E).
5. **Sequencing Ions with Base Loss (*s-B*)**: sequencing ions from which a neutral or charged sub-nucleotide fragment has detached (Figure 3F).

Further, sequencing ions are categorized into nine types, namely *a, b, c, d, a-B, w, x, y*, and *z*^22,23^, as illustrated in Figure 3. This figure provides a comprehensive visual reference for their annotation and interpretation. Users can enable or disable any ion types through the user interface of RNabel, which provides customization options for complex datasets or specific experimental requirements.

**Figure 3.**
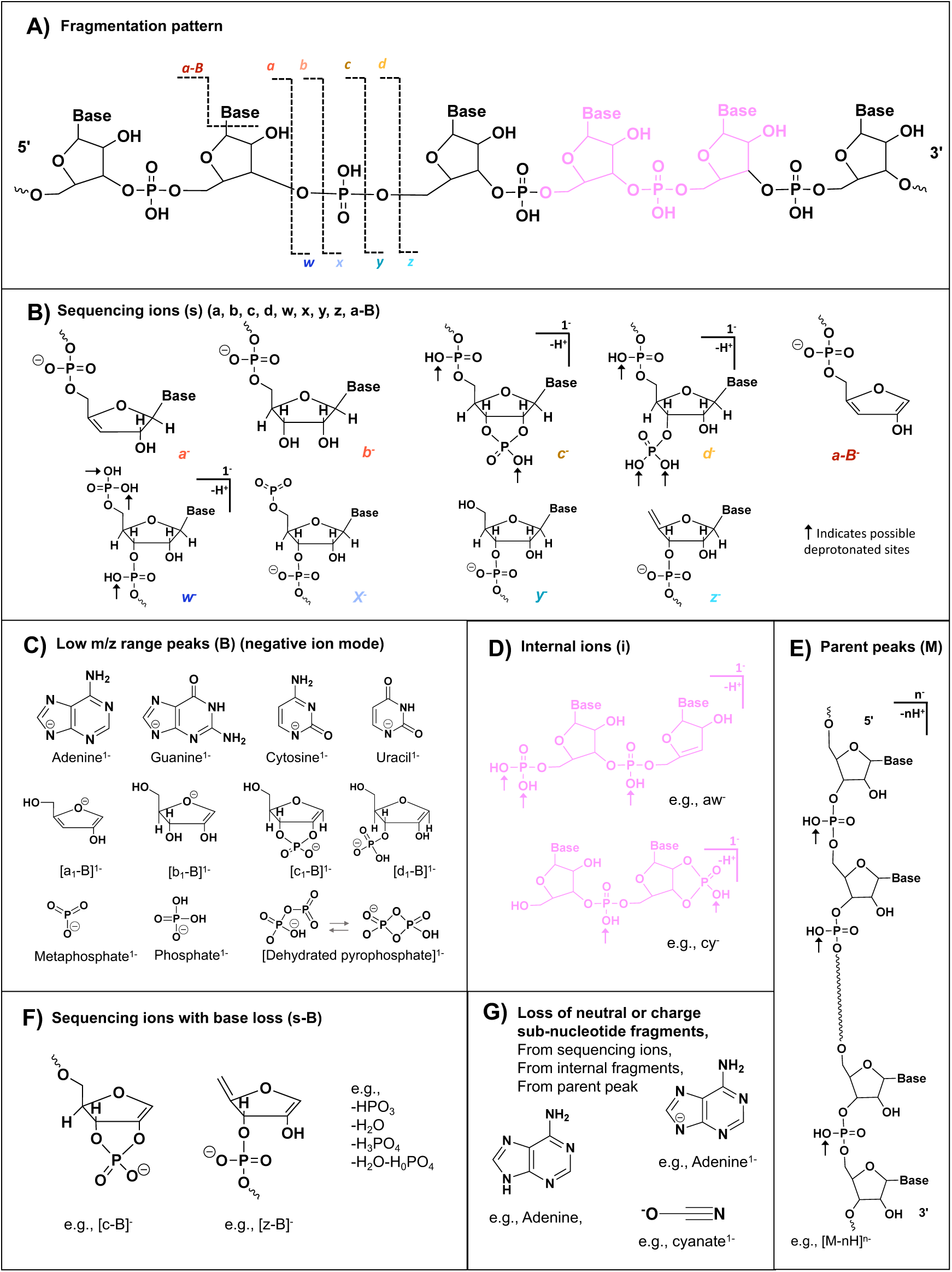
Theoretical fragment ions of RNA—chemical structures and classification in RNabel. A) Fragmentation sites and corresponding ion types. Two neighboring nucleosides and the phosphoryl group in between are colored pink to illustrate origin of internal ions (D). (B-E) Proposed chemical structures of different categories of fragment ions. Arrows indicate possible deprotonation sites. All ion types are incorporated into the RNabel workflow and mapped to one-letter codes for spectral labeling (see Table 1). Two structural isomers of dehydrated pyrophosphate ion (HP₂O₆⁻) ^42,43^ are shown (C). In addition to the predefined ion types shown here, RNabel also supports user-defined sub-nucleotide fragments, which can be applied to sequencing, internal, or parent ions (see Figure 4).

### Evaluation of ion-type abundance

Certain 5′ and 3′ fragment ions—specifically *a/z, b/y, c/x,* and *d/w* of the same nucleotide composition—can be isobaric, thereby confounding ion type determination. This is particularly common for RNA of palindromic sequences (as illustrated in Figure S2A).

To resolve such ambiguity, we designed a 20-nt RNA oligo whose first nucleotide is 2′-O-methylated A (Am). The 14.016 Da mass addition introduced to the 5′ end is expected to impart distinct molecular compositions for all fragment ions. Theoretical calculations of fragment m/z values confirmed this prediction; across experimentally relevant precursor charge states (z = 3^-^–7^-^) no more than four pairs of isobaric fragment ions occurred by chance (Figure S2B-C). In contrast, a 20-nt unmodified palindromic RNA may expect 26–44 pairs of isobaric fragment ions (±20 ppm) for precursor charge states of 3^-^–7^-^ (Figure S2C), which is in keeping with a previous study of shorter RNA oligos^34^.

HCD of this 2′-O-methylated oligonucleotide AmAAUCGAAAGCUAGCACAAA showed that that a/a-B, w, c, and y ions have the highest intensity and are also abundant (Figure S2D-E), confirming previous studies of collision-induced fragmentation of unmodified RNA oligonucleotides. Therefore, a/a-B, w, c, and y ion series are placed in the highest priority tier.

Internal ion formation requires the phosphodiester backbone to cleave twice, which is less probable under the conditions used, because they are optimized to enhance sequencing ions. Therefore, if a sequencing ion (Figure 3B) and an internal ion (Figure 3D) can both explain an experimental peak, the sequencing ion takes priority in RNabel.

Additionally, internal ions derived exclusively from backbone cleavage (Figure 3D) take priority over s-B ions (Figure 3F), because phosphodiester bond cleavage is energetically more favorable than glycosidic bond cleavage.

In sum, RNabel follows a four-tier priority rank, as shown below, to assign fragment ions.

First priority: a-B, a, w, c, y, M, and B ions.
Second priority: b, d, x, and z ions.
Third priority: i (internal) ions.
Fourth priority: s-B ions.

By default, RNabel includes only the first-priority ions in the annotation. Users have the option to enable second-, third-, or fourth-priority ions, or combination of them. Additionally, individual ion types within the same priority group can be enabled or disabled independently.

It should be noted that this priority hierarchy was established based on the fragmentation behavior of unmodified RNA oligonucleotides under collision-induced activation (HCD/CID). This may not be applicable to all modification types or fragmentation methods. For example, alternative fragmentation methods (e.g., NETD^37^, UVPD^38^, a-EPD^38^, IRMPD^39^) and backbone modifications such as phosphorothioate substitutions can alter the relative abundance of different ion series. To accommodate such cases, RNabel provides a dedicated Ion Priority Config panel (accessible via the Tools menu) for users to define custom ion priority schemes.

### Customized Annotation

Users can define custom nucleotides, modifications, 5′ and 3′ terminal groups, neutral losses, and charged sub-nucleotide fragments (Figure 4). This allows researchers to tailor the annotation process to accommodate novel or uncommon RNA/DNA modifications.

**Figure 4:**
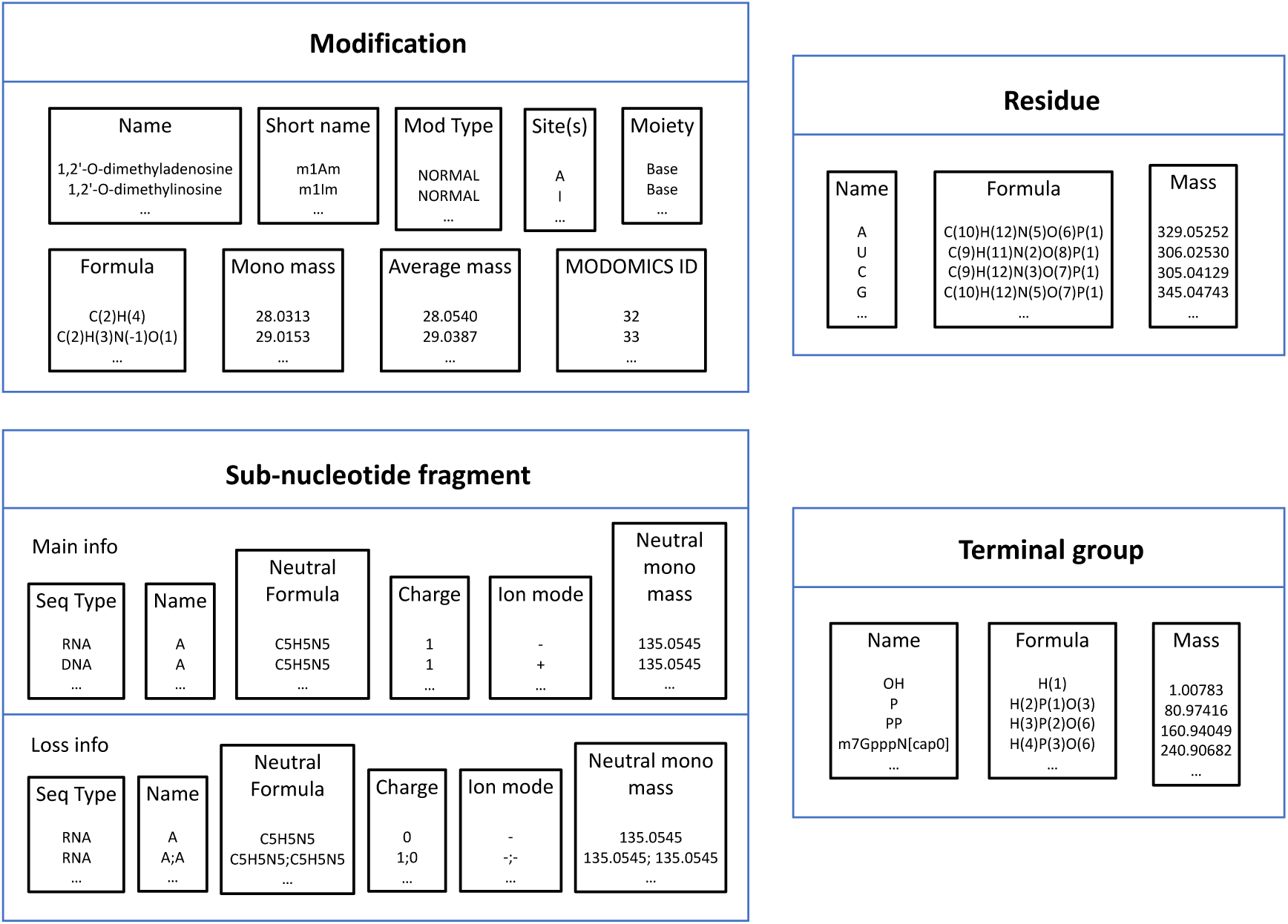
Customization options in RNabel. RNabel users have the option to create customized annotation of RNA or single-stranded DNA spectra. Users can define new modifications by specifying the full name, short name (used for visualization on the annotated spectra canvas), chemical formula, affected moiety (base, ribose, or phosphate), mass shift (monoisotopic mass and average mass), and the MODOMICS ID (if available, or −1). This allows the annotation of novel or uncommon RNA modifications. Users can also define custom nucleotide residues by providing their name, chemical formula, and monoisotopic mass, and have them incorporated into the analysis. A sub-nucleotide fragment is characterized by two components: main information (“main info”) and loss information (“loss info”). Both components include the sequence type (RNA/DNA), name, neutral chemical formula (required for both charged and neutral losses), charge state, and ion mode. The “loss info” allows multiple losses to be defined for a single sub-nucleotide fragment, with each loss separated by a semicolon (;). These definitions are applied to fragment ions to annotate characteristic neutral or charged sub-nucleotide fragments observed in RNA MS/MS spectra. Lastly, users can define terminal groups at the 5′ and 3′ ends by specifying their name, chemical formula and monoisotopic mass.

#### RNA Modification

RNabel supports all RNA modifications listed in the MODOMICS database (https://genesilico.pl/modomics/). Users can also define new RNA modifications by specifying the following parameters: **Name** (full modification name); **Short Name** (abbreviated name of a modification to appear in annotated spectra); **Mod Type** (modification type, including Oligo 5-term (occurs only at the 5′ terminal of an oligo), Oligo 3-term, RNA 5-term (occurs only at the 5′ terminal of an RNA molecule), RNA 3-term, NORMAL (modification position not restricted to terminal positions)); **Sites** (nucleotide to receive the modification, options typically include A, U, C, G, or N, of which N is any nucleotide); **Moiety** (component of a nucleotide being modified, such as ribose, base, phosphate, or undefined); **Formula** (chemical formula representing the mass shift caused by the modification); **Monoisotopic Mass** (monoisotopic mass of the modification); **Average Mass** (average mass); **MODOMICS Database ID** (unique ID from the MODOMICS database, or −1 for novel modifications). Figure S5 illustrates the graphical user interface and step-by-step instructions for defining new RNA modifications.

#### Nucleotide Residue

RNabel allows users to define custom nucleotide residues, providing flexibility beyond the four standard RNA and DNA nucleotides. This feature significantly enhances RNabel’s scalability, making it ideal for analyzing non-canonical nucleotide bases. Users can define a custom nucleotide residue by specifying the following: **Name** (name of the nucleotide residue); **Formula** (chemical formula); **Mass** (monoisotopic mass); **Neutral Base Formula** (chemical formula of the nucleobase); **Neutral Base MonoMass** (monoisotopic mass of the nucleobase). Figure S6 illustrates the graphical user interface and the step-by-step instructions for defining new nucleotides.

For example, phosphorothioate (PS) RNA, where one or more backbone phosphates are modified with a sulfur atom instead of the standard non-bridging oxygen, is a common modification due to its ease of synthesis and pharmacokinetic benefits. To define a custom PS adenosine in RNabel, the user would specify:

Name: A*
Formula: C(10)H(12)N(5)O(5)P(1)S(1)
Mass: 345.0296767
Neutral Base Formula: C(5)H(5)N(5)
Neutral Base Monomass: 135.05449518285

#### 5′ and 3′ terminal group

Users can define own 5′ or 3′ terminal groups by providing **Name** (name of the terminal group), **Formula** (chemical formula), and **Mass** (monoisotopic mass). With this extended feature, RNabel can accommodate a wide range of terminal modifications. Figure S7 provides the graphical user interface and step-by-step instructions for defining new terminal groups.

Users define custom 5′ or 3′ terminal groups by providing the following key parameters: **Name** (name of the terminal group); **Formula** (chemical formula); **Mass** (monoisotopic mass).

#### Loss of Neutral or Charged Sub-Nucleotide Fragment

Evidence in MS/MS indicates that a sub-nucleotide fragment—as either a neutral or a charged group—may come off an RNA precursor ion or a standard fragment ion (*a, b, c, d, a-B, w, x, y,* and *z*). RNabel divides these outcome of such events into three categories: (1) low-*m/z* sub-nucleotide fragment ions including base, ribose (e.g., a₁–B₁⁻), phosphorated ribose (e.g., d₁–B₁⁻), phosphate, pyrophosphate, and cyanate (Figure 3C and G), (2) precursor ions from which a neutral or charged (or combinations of both) sub-nucleotide fragment has broken off, (3) sequencing or internal ions from which with a neutral or charged (or combinations of both) sub-nucleotide fragment has broken off.

A custom neutral/charged loss in RNabel is defined by information about neutral/charged loss and information about the source ion that generates such a loss.

Information about the neutral/charged loss includes: **Loss Seq Type** (DNA or RNA); **Loss Ion Mode** (+ or −); **Loss Name** (name of the loss group being defined); **Loss Charge** (charge state of the lost group, a positive or negative integer number for charged loss, 0 for neutral loss); **Loss Neutral Formula** (formula of the loss group expressed as a neutral); **Loss Num** (an integer number indicating how many copies are lost, e.g. 2 for the loss of 2 phosphate groups); **Loss Ion Mass** (a negative value indicating the total mass that is lost, e.g. −195.95379 for the loss of 2 phosphate groups).

Information about the source ion includes (input needed only for low m/z range peaks): **Main Seq Type** (DNA or RNA); **Main Ion Mode** (“+” or “−”); **Main Name** (name of the main fragment); **Main Charge** (charge state of the main fragment, a positive integer number); **Main Neutral Formula** (formula of the source ion expressed as a neutral); **Main Neutral Mass** (neutral mass of the source ion).

RNabel also supports combinations of two or more neutral/charged losses to help dissect complex fragmentation patterns. For example, to inspect a simultaneous loss of a phosphoric acid and a water, users can define a custom sub-nucleotide fragment by specifying both groups and separating them with a semicolon. In this case, the parameters would be:

- Loss Name: H_3_PO_4_;H_2_O
- Loss Neutral Formula: H_3_PO_4_;H_2_O
- Loss Charge: 0;0
- Loss Num: 1;1 (indicating that both losses occur once)
- Loss Ion Mass: −97.97690;-18.01056

Since this loss belongs to the **Parent Sub-nucleotide Fragment** category, the main peak is permitted to be any of the precursors, so the Main Information is not required and left empty.

### Isotopic Distribution Matching (IDM)

Compared to an average amino acid residue^40^, an average mononucleotide of A, U, C, G has almost twice as many carbon atoms to incorporate the naturally occurring heavy isotope ^13^C. Consequently, as the length of an RNA oligo increases, the monoisotopic peak M quickly diminishes while the isotopic peaks grow. Simulation shows that once an RNA is more than 8 nt long (or beyond ∼3900 Da), the monoisotopic peak is no longer the tallest peak—namely, the base peak—in the isotopic distribution (Figure 8A). For a 32-nt RNA, the monoisotopic peak accounts for only 1% of the isotopic distribution. Shown in Figure 8B-E, as an RNA increases from 4 nt to 32 nt, the base peak shifts gradually from the monoisotopic peak to the 4^th^ isotopic peak. Most available software tools except Aom^2^S^32^ annotate only the monoisotopic peaks. Clearly, this needs improvement.

In RNabel we implemented the Isotopic Distribution Matching (IDM) algorithm (Figure S12). For each candidate fragment, IDM generates a theoretical isotopic distribution and calculates a cosine similarity score between the theoretical and the experimental isotopic distribution.

Next, we examined the projections of truly or falsely matched isotopic distributions on the cosine similarity score and on mass deviation (Figure S13). Using a data-adaptive 3-component Gaussian Mixture Model (GMM), we evaluated the IDM matching results of Dataset #2, which contains 333 CID/HCD spectra of 19 RNA oligos of 16–32 nt. As shown in Figure S13A, this analysis revealed three clusters, recognized as ‘True matches’, ‘Gray zone’, and ‘Noise’. True matches are high in cosine similarity (97% scored 0.8–1.0) and low in mass deviation (100% within ± 5 ppm). Gray zone and Noise matches displayed low cosine similarity (< 0.8 and < 0.4, respectively) and large mass deviations (41.7%–53.2% ≥ 5 ppm). As such, cosine similarity ≥ 0.8 and mass deviation within ± 5 ppm are set threshold values for IDM matching.

It is important to emphasize that the cosine similarity boundary derived here is data-dependent, and it could change with isolation window, MS2 resolution and mass accuracy, and instrument type, etc. Therefore, we strongly recommend that users apply this GMM clustering approach to own data to determine appropriate threshold values.

### Graphical User Interface

RNabel features an intuitive graphical user interface (GUI) to streamline the annotation process. The GUI is organized into four main sections: the **Toolbar Area**, where users can load spectrum files, define new nucleotides, modifications, terminal groups, and custom sub-nucleotide fragments; the **Spectrum Display Area**, which shows annotated or unannotated MS2 spectra with color-coded ions for easy identification, and offers customization options for ion colors; the **Statistical Parameter Display Area**, which presents key statistical metrics such as sequence and position coverage, and interpretation rates for different ion types; and the **User Input Parameter Area**, where users enter sequence information (e.g., RNA/DNA, nucleotide sequence, terminal groups, modifications, and charge state of the parent ion). For a visual representation of the GUI, see Figure 5.

**Figure 5.**
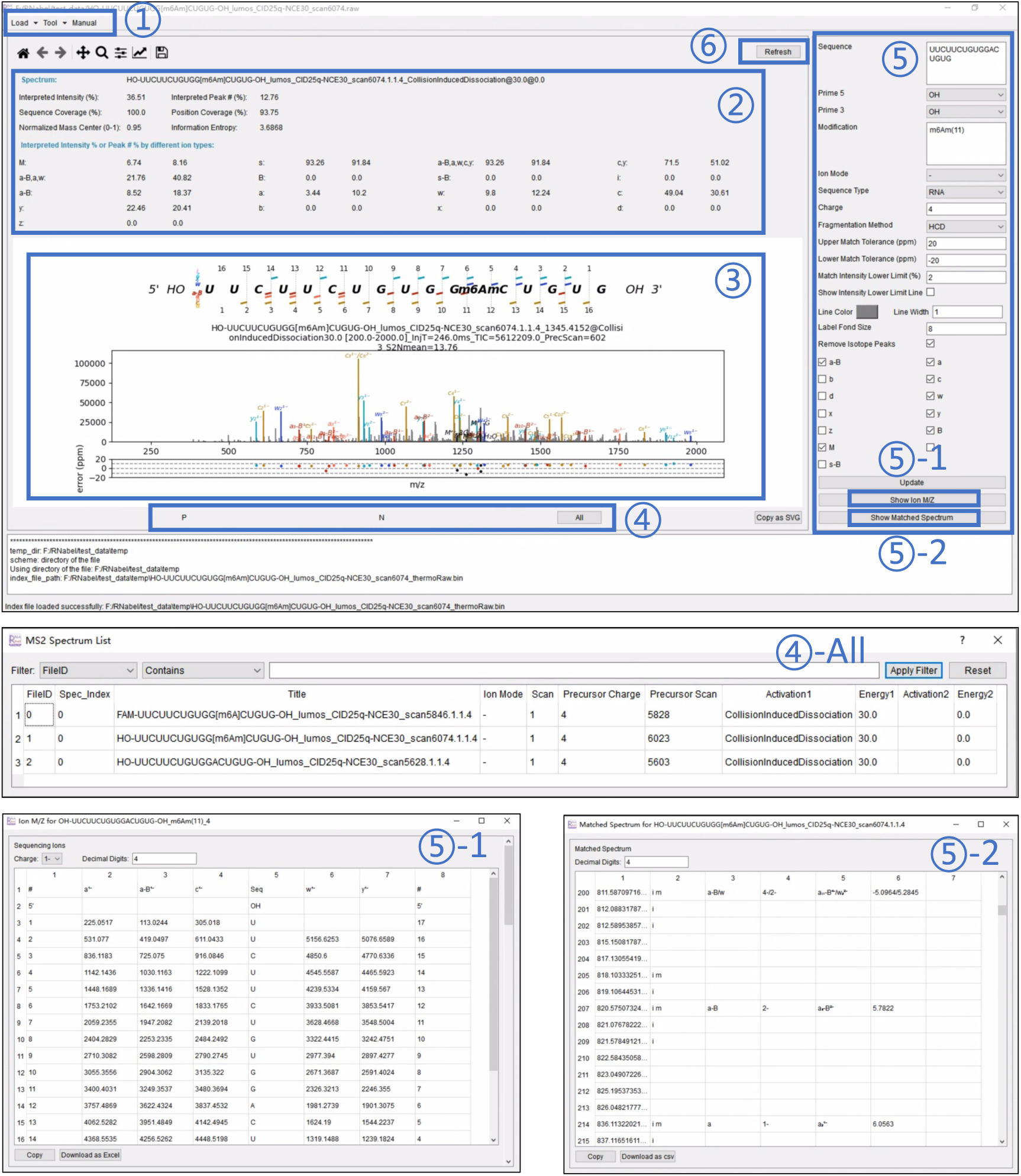
The graphical user interface of RNabel. There are six functional regions in the graphical user interface of RNabel: **①** Input and tool buttons: Allow the input of tandem MS data of RNA or DNA in mgf, msConvert2 text, Thermo raw, or RNabel batch txt formats, as well as the definition of new modifications, sub-nucleotide fragments, RNA/DNA nucleotides, and 5′/3′ terminal groups. **②** Statistical indicators region: Summarizes sequence and position coverage metrics and the interpretation rates of each type of sequencing ion in the annotated spectrum. **③** Matched spectra region: Displays the fragmentation map, the annotated tandem MS spectrum, and an error plot of the selected spectrum. These plots can be downloaded in vector graphics format. **④** Navigation buttons and spectrum list window: The “P” and “N” buttons navigate to the previous or next spectrum in the dataset, respectively. **④-All** Clicking the “All” button opens a window listing all spectra in the dataset; selecting a row directly navigates to the corresponding spectrum. **⑤** Sequence information window: In this window, users enter nucleic acid sequence, 5’ and 3’ termini, modifications, ion mode, and error tolerance. RNabel supports all modifications cataloged in the MODOMICS database. The update button calculates the theoretical fragment ions according to the sequence information and annotates the loaded spectrum. **⑤-1** Theoretical ion m/z table: Displays the m/z of theoretical ions in tabular format **⑤-2** Matched Spectrum Table: Displays matched spectrum details in tabular format. **⑥** The Refresh Button: Resets the spectrum coordinate axes to the default view.

### File Format Support

RNabel supports input files in the following formats: MGF, Thermo RAW, msConvert text, and RNabel batch files. Detailed specifications for the RNabel batch file are available in Table S2.

### Other Features

RNabel offers several practical features designed to enhance usability and improve the efficiency of spectrum analysis, as follows.

1) supports annotation of MS/MS spectra of single-stranded DNA oligos. 2) allows users to adjust font size, and ion type colors, zoom in and out of spectra, and download theoretical ion masses in Excel format. 3) easy to browse through MS/MS spectra via a window listing loaded spectra. 4) provides statistics for evaluating the quality of annotation, as detailed in Methods. 5) easy access to documentation via the Help menu.

### Display of Annotated MS/MS Spectrum

RNabel provides high-quality annotations for RNA tandem MS spectra, allowing users to visualize and interpret complex fragmentation patterns. An annotated MS/MS spectrum of RNA consists of a fragmentation map, the matched spectrum itself, and an error plot. The **fragmentation map** illustrates the matching of fragment ions along the RNA backbone, highlighting sequence fragments at specific positions. This facilitates understanding of the fragmentation pattern along the RNA chain. Below the fragmentation map, the **matched spectrum** displays the experimental peaks that have been assigned to theoretical ions, with distinct colors denoting various ion types. Finally, the **error plot** provides a graphical representation of the mass differences between the theoretical and experimental peaks, sharing the same x-axis as the matched spectrum.

Figures 6 and 7 display multiple RNabel-annotated MS/MS spectra of modified and unmodified RNA oligos in both negative and positive ion modes, with different 5′ terminal groups, modifications and fragmentation methods (CID and HCD). This demonstrates RNabel’s versatility and effectiveness in MS/MS spectrum annotation for nucleic acid analysis.

**Figure 6.**
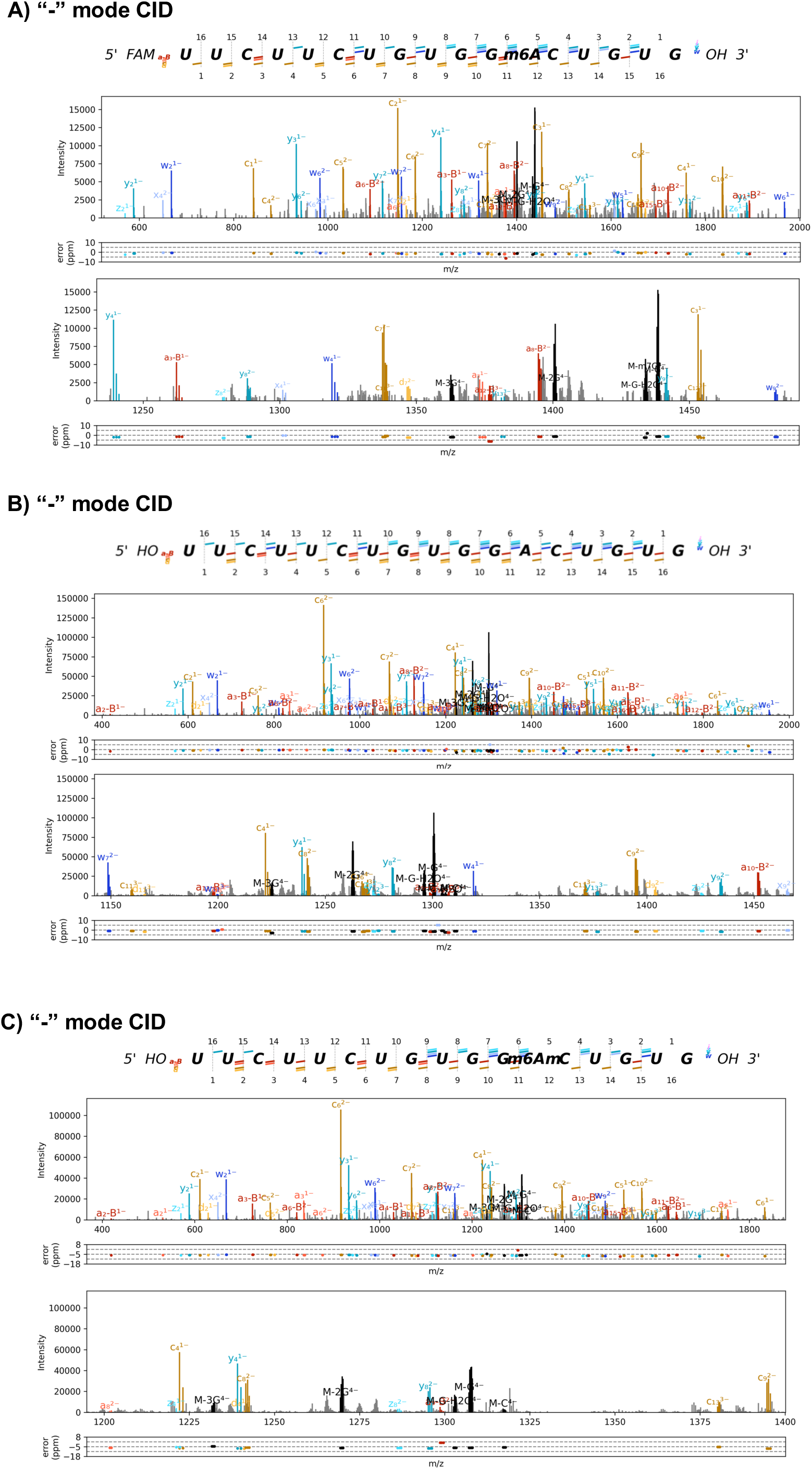
Annotated negative ion mode MS/MS spectra of modified and unmodified RNA oligonucleotides. Panel (A) shows the annotated spectrum for an RNA sequence modified with m6A and featuring a FAM group at the 5′ terminus. Panel (B) displays the spectrum for an unmodified RNA sequence with a hydroxyl group at the 5′ end. Panel (C) illustrates the annotated spectrum for an RNA sequence containing the m6Am modification, also with a hydroxyl group at the 5′ end. For each panel, the top plot displays the full MS/MS spectrum, while the bottom plot provides a zoomed-in view of a specific *m/z* range to zoom in the detailed fragment ion annotations. All spectra were acquired using collision-induced dissociation (CID) at a normalized collision energy (NCE) of 30. Detailed segment-by-segment zoomed-in views for each annotated spectrum are provided in Figures S9, S10, and S11 for Panels (A), (B), and (C), respectively. These examples demonstrate RNabel’s capability to accurately annotate both modified and unmodified RNA oligos with different terminal groups, showcasing its versatility and effectiveness in RNA mass spectrometry analysis.

**Figure 7.**
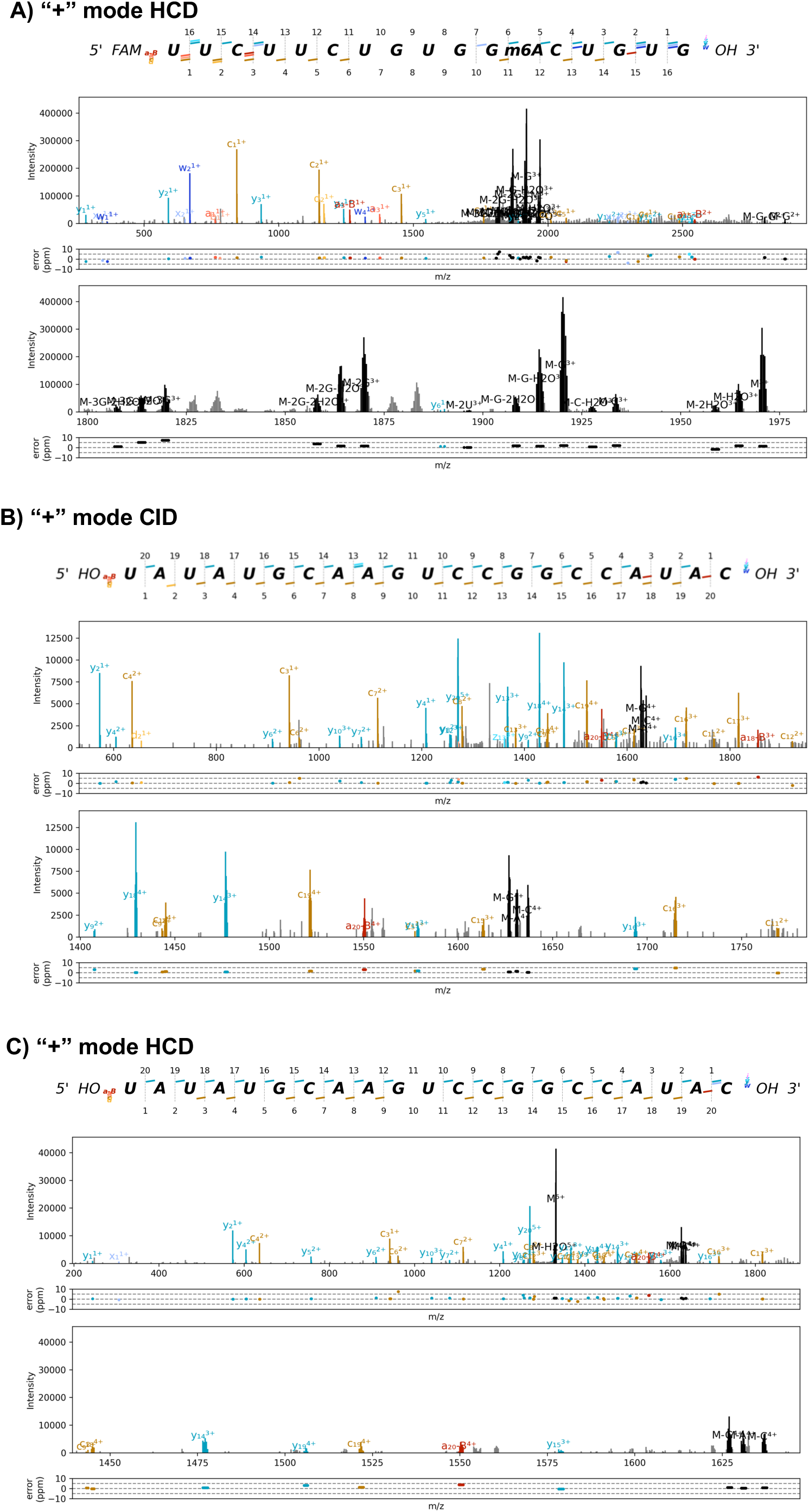
Annotated positive ion mode MS/MS spectra of modified and unmodified RNA oligonucleotides acquired under different fragmentation conditions. Panel (A) shows the annotated spectrum for the m⁶A-modified RNA oligonucleotide (FAM-m6A-OH, 5ʹ-FAM-UUCUUCUGUGG[m⁶A]CUGUG-OH-3ʹ) with a FAM group at the 5ʹ terminus, acquired using higher-energy collisional dissociation (HCD). Panel (B) displays the spectrum for an unmodified RNA oligonucleotide (5ʹ-HO-UAUAUGCAAGUCCGGCCAUAC-OH-3ʹ, 21-mer) acquired using collision-induced dissociation (CID). Panel (C) presents the spectrum for the same unmodified RNA sequence as in Panel (B), acquired using HCD. For each panel, the top plot displays the full MS/MS spectrum, while the bottom plot provides a zoomed-in view of a specific *m/z* range to highlight the detailed fragment ion assignments. These examples demonstrate RNabel’s capability to annotate RNA oligonucleotides in positive ion mode, complementing the negative ion mode results shown in Figure 6.

**Figure 8:**
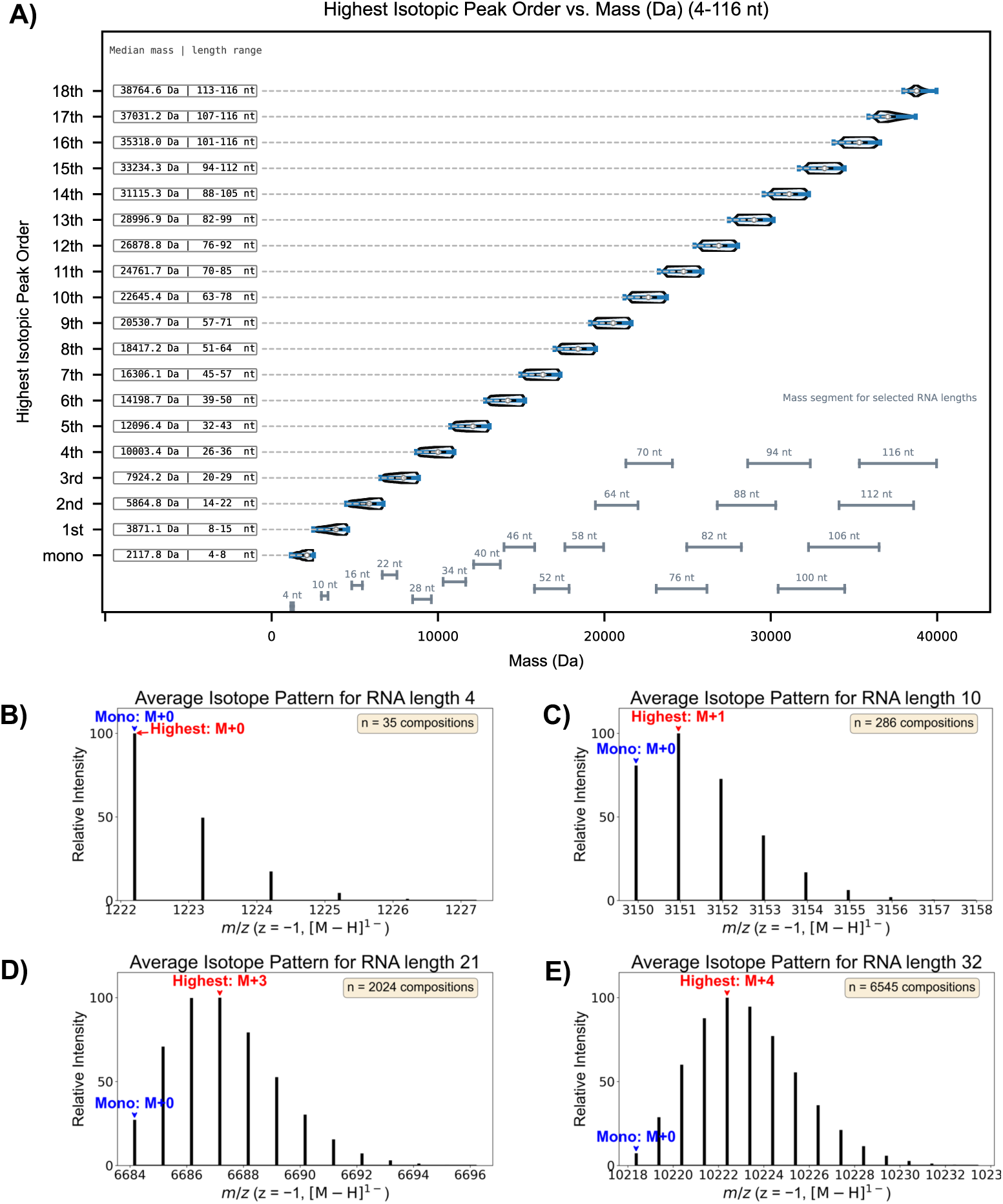
Simulation of isotopic distribution patterns of RNA oligonucleotides. (A) The base peak (peak of the highest intensity) shifts from the monoisotopic peak (M+0) to the 18^th^ isotopic peak as the length of RNA increases from 4 nt to 116 nt, or from just above 1000 Da to 40,000 Da. Each of the horizontal dashed lines at the bottom denotes the theoretical mass range of an oligonucleotide of the indicated length. (B-E) Simulated average isotopic distributions of4-, 10-, 21-, and 32-nt RNA oligos.

## Discussion

This study introduces RNabel, a versatile tool for tandem mass spectrum annotation of nucleic acids. Designed with both novice and expert users in mind, RNabel simplifies the annotation task by offering a friendly graphical user interface (GUI) with robust functionalities.

RNabel stands apart from existing tools by supporting users to define custom RNA modifications, nucleotide residues, terminal groups, neutral losses, charged sub-nucleotide fragments detached from backbone fragment ions, and combination of the last two groups (e.g., simultaneous loss of a water molecule and a dihydrogen phosphate ion from an a-B ion). RNabel facilitates customization by presenting an intuitive GUI. This flexibility enables annotation of oligonucleotides bearing non-standard modifications. Also, it provides a platform for defining diagnostic ions to distinguish isobaric species. For example, it is reported that the stable C–C glycosidic bond of pseudouridine leads to unique MS/MS fragments of m/z 164 and m/z 165, which are not produced by uridine because it bears a more labile C-N glycosidic bond^41^. Such customization empowers users to explore novel RNA modifications and complex nucleic acid chains.

To meet the demand of high-throughput analysis of RNA samples, RNabel supports batch annotation. RNabel’s multi-level indexing strategy is memory-efficient, enabling simultaneous processing of multiple files and large datasets without system overload. This utility is further enhanced by a set of integrated statistical metrics for evaluating the quality of MS/MS annotation.

Overall, RNabel offers researchers a powerful, scalable solution for analysis of RNA MS/MS spectra. Future developments will focus on: (1) enhanced support for longer RNA oligonucleotides; (2) integration of error-aware scoring functions to improve annotation accuracy for internal fragment ions; (3) implementation of de novo sequencing capabilities to enable sequence determination of unknown RNA species; and (4) extension to new application domains, such as RNA-protein crosslinking analysis.

## Acknowledgments

The authors would like to thank Dr. Si-Min He, Dr. Chi Hao, and other members of the pFind group, ICT, CAS for guidance to programming. The authors gratefully acknowledge grant support from the National Science Foundation of China (Grant No. 32571674) and NIBS, Beijing, whose intramural grants are funded by the Ministry of Science and Technology of China, the municipal government of Beijing, and Tsinghua University.

## Data Availability

The raw RNA MS/MS spectra supporting this study have been deposited in the MassIVE repository under accession number MSV000100052. The dataset can be accessed via FTP at ftp://MSV000100052@massive-ftp.ucsd.edu using the username MSV000100052 and password m8C7k8suMTqwVQiH. The RNabel software is available at https://github.com/songge1111/RNabel/releases.

## Notes

ftp://

https://github.com/songge1111/RNabel/releases

### Competing Interest Statement

The authors have declared no competing interest.

