## Supplemental file for "RNabel—A Standalone Software Tool for Annotating Tandem Mass Spectra of Modified Ribonucleic Acids"

**Song et al.**

**Contents**

Supplementary Figure 1: An annotated RNA MS/MS spectrum generated by a third-party software tool

Supplementary Figure 2: Resolving isobaric fragment ions using an RNA oligo containing a 2'-O-Methylated ribose in the first nucleotide

Supplementary Figure 3: Efficient multi-file loading and indexing mechanism in RNabel

Supplementary Figure 4: Workflow for batch spectrum annotation in RNabel

Supplementary Figure 5: Customization GUI for RNA Modifications in RNabel

Supplementary Figure 6: Customization GUI for Defining New Residue Types of RNA/DNA in RNabel

Supplementary Figure 7: Customization GUI for Defining New Terminal Groups in RNabel

Supplementary Figure 8: Customization GUI for Defining New Sub-nucleotide fragments in RNabel

Supplementary Figure 9: Detailed segment-by-segment zoomed-in views of the annotated MS/MS spectrum for the m⁶A-modified RNA oligonucleotide (FAM-m6A-OH, 5ʹ-FAM-UUCUUCUGUGG[m⁶A]CUGUG-OH-3ʹ), corresponding to the spectrum shown in Figure 6A

Supplementary Figure 10: Detailed segment-by-segment zoomed-in views of the annotated MS/MS spectrum for the unmodified RNA oligonucleotide (HO-A-OH, 5ʹ-HO-UUCUUCUGUGGACUGUG-OH-3ʹ), corresponding to the spectrum shown in Figure 6B

Supplementary Figure 11: Detailed segment-by-segment zoomed-in views of the annotated MS/MS spectrum for the m⁶Am-modified RNA oligonucleotide (HO-m6Am-OH, 5ʹ-HO-UUCUUCUGUGG[m⁶Am]CUGUG-OH-3ʹ), corresponding to the spectrum shown in Figure 6C

Supplementary Figure 12: Workflow of the isotope distribution matching (IDM) algorithm

Supplementary Figure 13: Unsupervised Gaussian Mixture Model (GMM) to determine the cosine similarity threshold

Supplementary Figure 14: Interactive mirror-plot interface for inspecting a match between two isotopic distributions

Supplementary Table 1: Statistical metrics calculated by RNabel to evaluate tandem MS spectrum annotation quality

Supplementary Table 2: Required and optional columns in RNabel batch input files

Supplementary Table 3: Synthetic RNA oligos used in the development of RNabel

Supplementary Table 4: Software comparison

**Supplementary Figures**

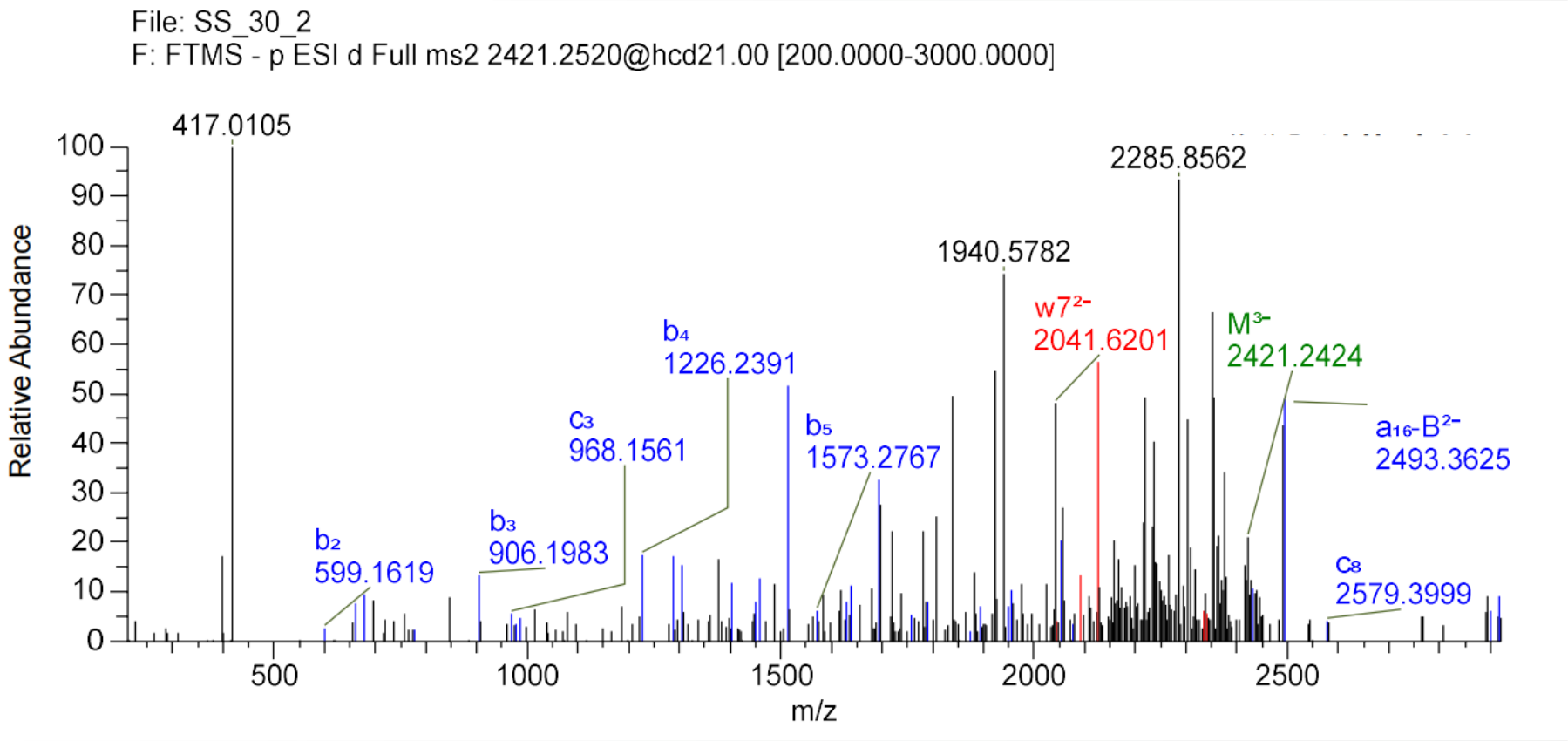

**Supplementary Figure S1: An RNA MS/MS spectrum of a 17-mer annotated by a third-party software tool.**

The sequence was 5ʹ-AmCmCfUmGfUmTdUmUmGmCmUmUmUmUmGmUm-3ʹ, where subscripts m, f, and d denote 2ʹ-O-methyl, 2ʹ-fluoro, and 2ʹ-deoxy modifications, respectively. Data were acquired on an Orbitrap mass spectrometer using higher-energy collisional dissociation (HCD) at a normalized collision energy of 21. Although some sequencing ions (b, c, w, and a−B series) provided sequence coverage, a considerable number of abundant peaks in the spectrum remained unannotated. These unexplained signals may originate from co-eluting species or atypical fragmentation pathways of the same precursor, potentially leading to ambiguity in sequence assignment.

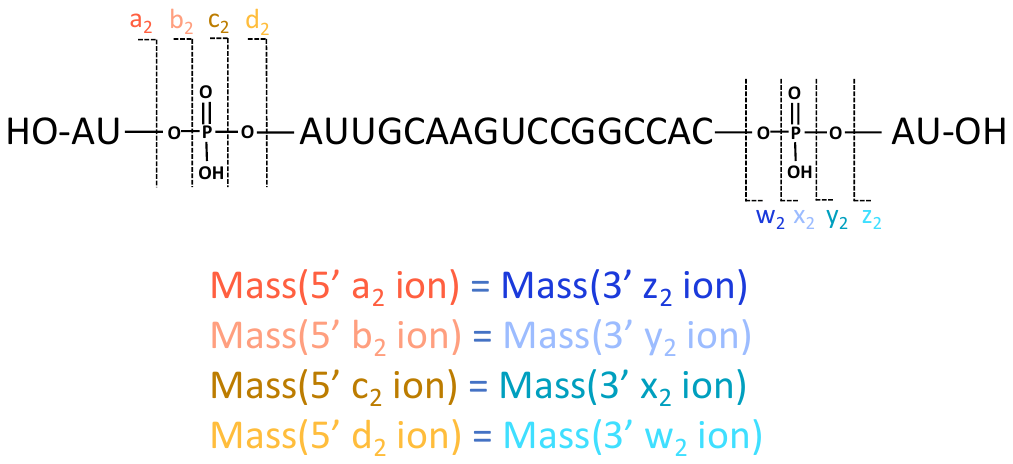

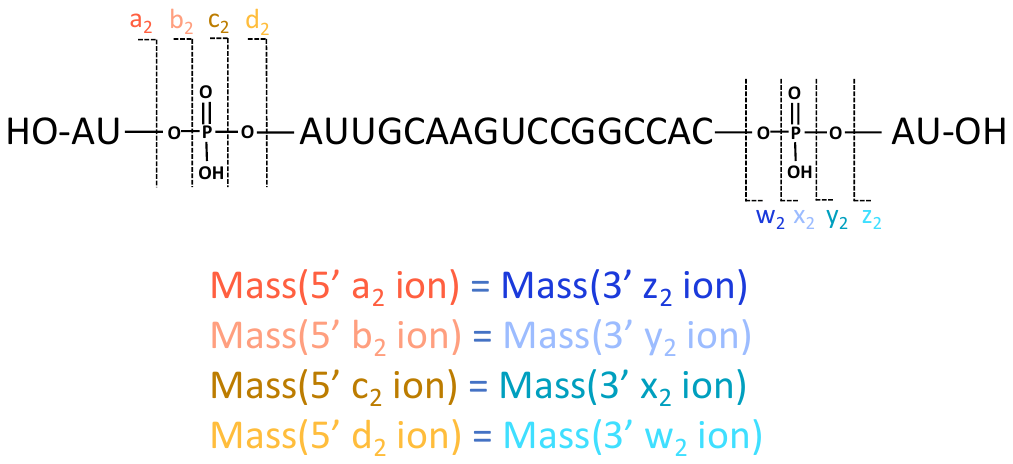

**A)**

**B)**

**C)**

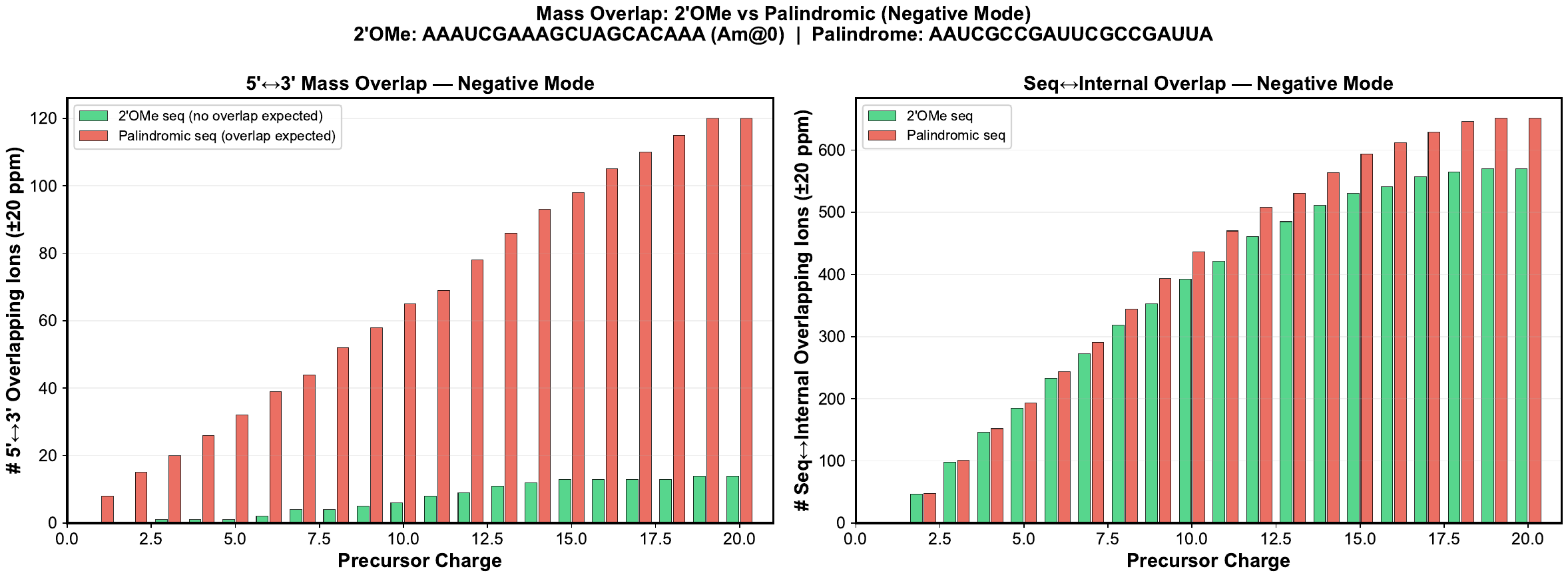

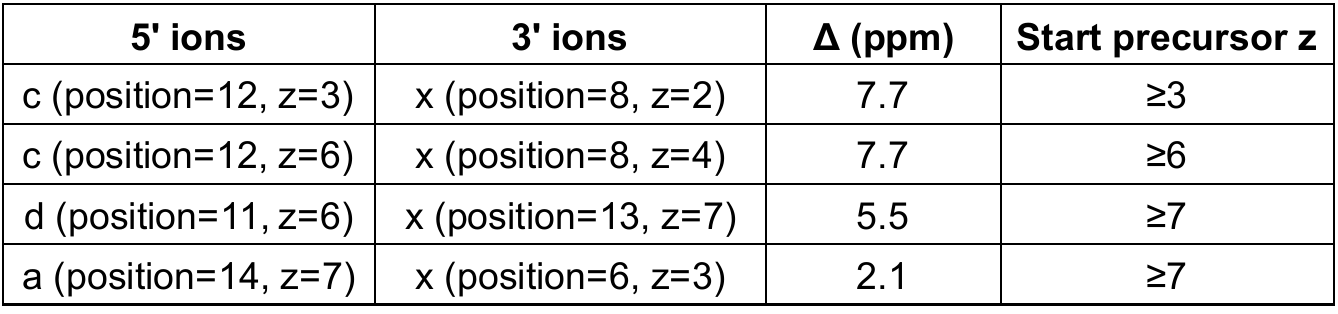

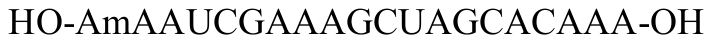

**D)**

**E)**

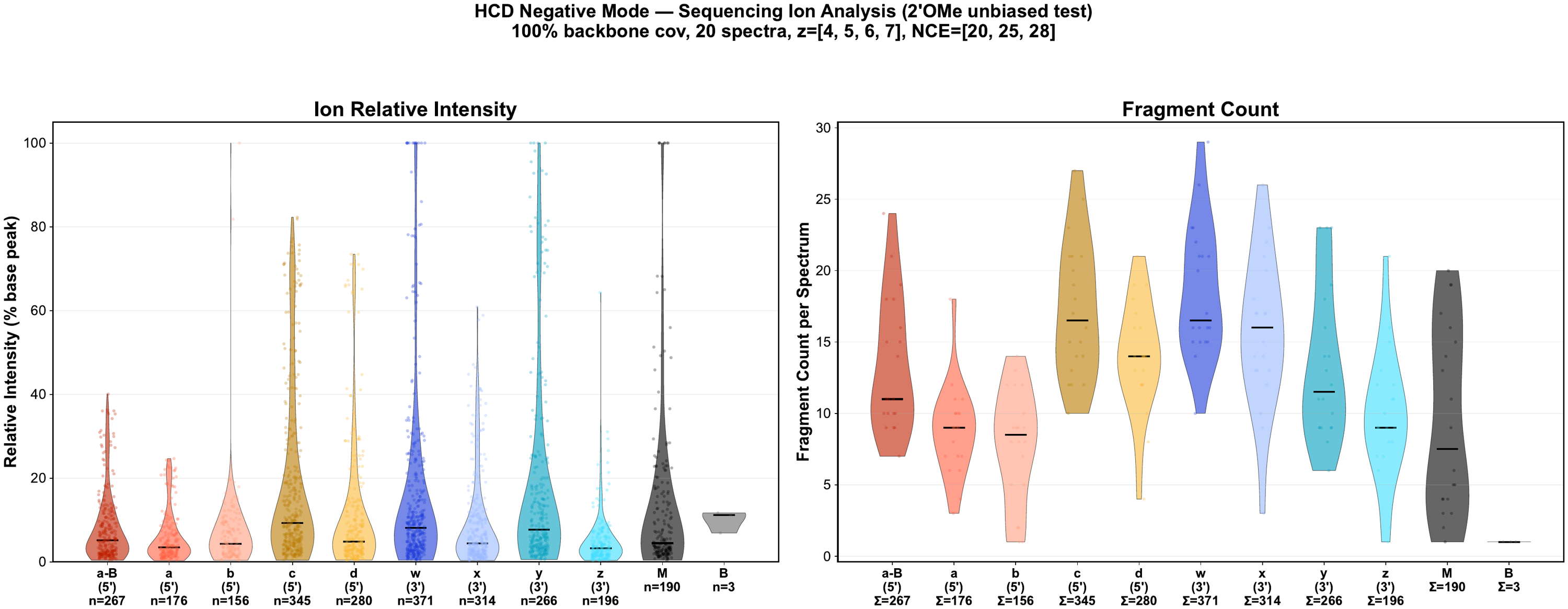

**Supplementary Figure S2: Resolving isobaric fragment ions using an RNA oligo containing a 2'-O-Methylated ribose in the first nucleotide.**

(A) Illustration of isobaric fragment ions using a palindromic RNA oligo. As shown, pairs of *a-* and *z*-ions, *b-* and *y*-ions, *c-* and *x-*ions, *d-* and *w-*ions are of the same composition and thus of the same m/z. (B) Based on the theoretical fragment ion m/z values calculated for the 2'-O-Methylated RNA (AmAAUCGAAAGCUAGCACAAA), only four pairs of isobaric ions may be expected from a precursor carrying 7 negative charges or less. (C) Number of possible isobaric ions pairs (or groups) (mass deviation ±20 ppm) calculated for the above methylated RNA (B) and a palindromic RNA (HO-AAUCGCCGAUUCGCCGAUUA-OH) under the indicated precursor charges state. (D-E) Relative intensity (D) and counts (E) of the different types of ions in an averaged HCD spectrum of the 2'-O-Methylated RNA above. A total of 20 spectra were analyzed; negative ion mode; z = 4–7; NCE = 20, 25, and 28; position coverage 100%.

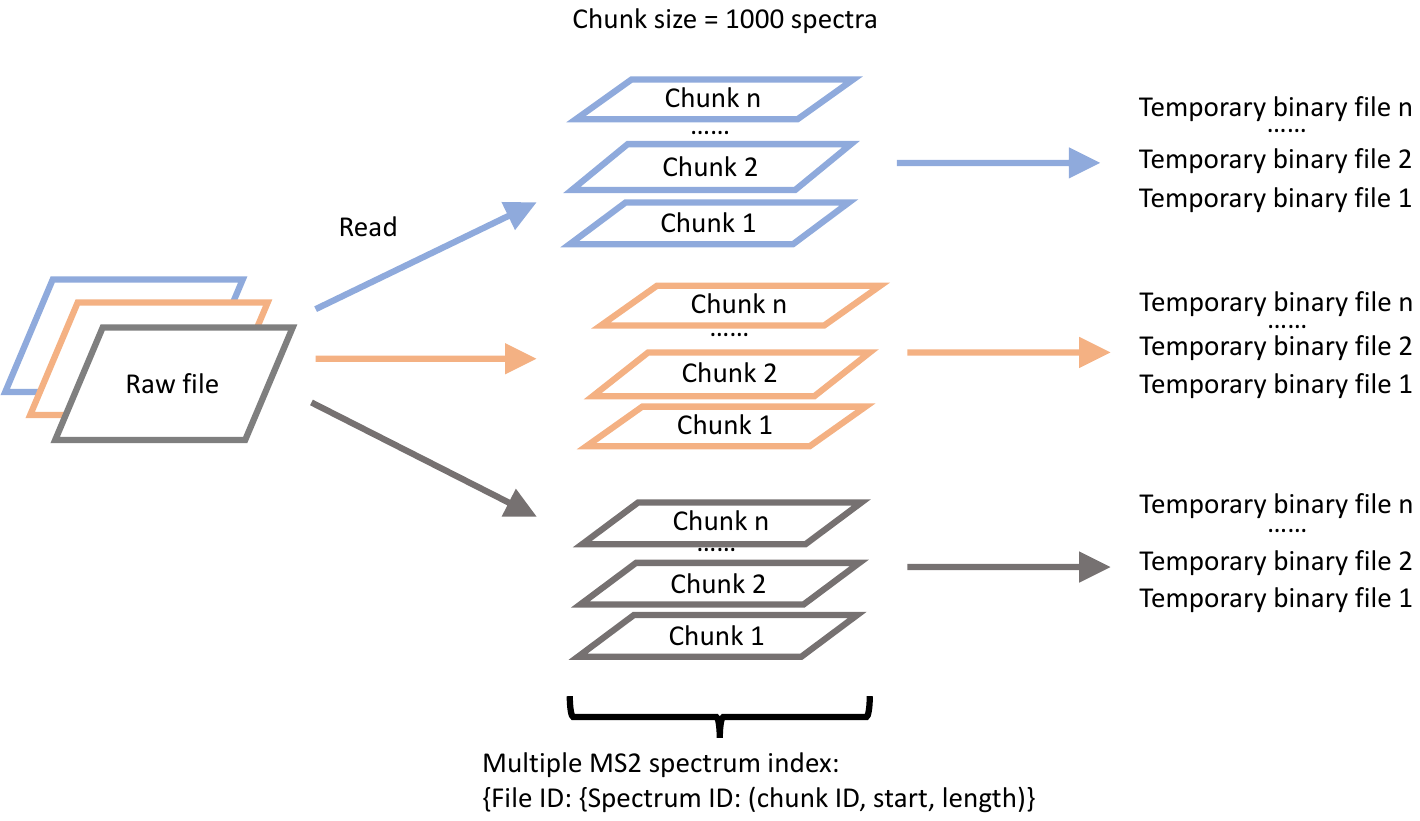

**Supplementary Figure S3: Efficient multi-file loading and indexing mechanism in RNabel**

RNabel employs an efficient multi-file loading strategy to handle large or multiple tandem MS data files without compromising performance. Each raw spectrum file is partitioned into independent chunks, which are indexed and loaded in parallel. This indexing mechanism allows for fast random access to individual spectra, minimizing memory consumption and improving responsiveness, especially when browsing large datasets. Each chunk can be retrieved on demand, enabling users to annotate spectra incrementally without reloading entire files. This design supports streamlined exploration and visualization of high-throughput RNA MS/MS data.

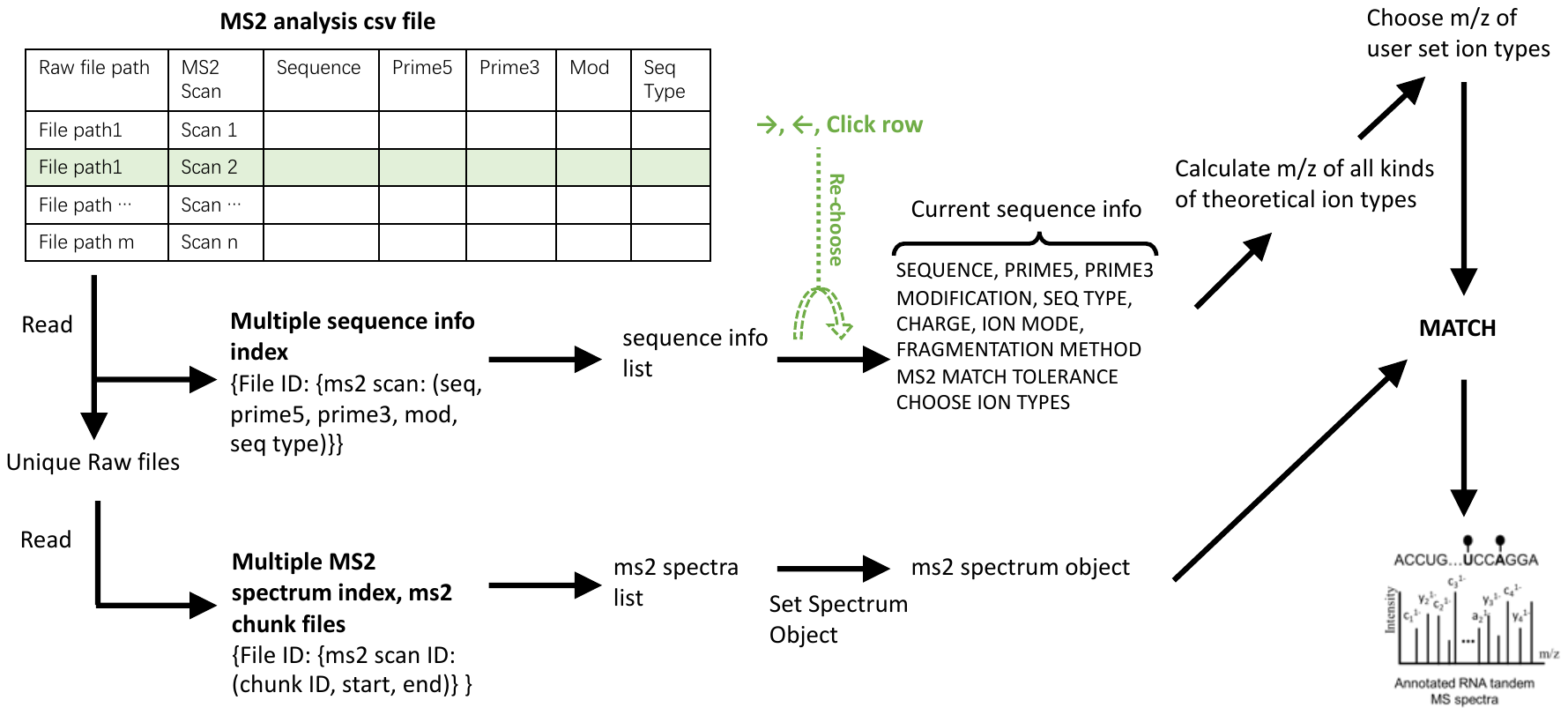

**Supplementary Figure S4: Workflow for batch spectrum annotation in RNabel.**

This figure illustrates the workflow for batch spectrum annotation in RNabel. Users first import a CSV file containing essential spectrum identity information, including the raw file path, MS2 scan number, RNA oligo sequence, 5′ and 3′ termini, modifications, and sequence type (RNA/DNA). Upon loading the batch file, RNabel automatically processes each entry and constructs an internal list of annotation tasks. Users can then navigate through the annotated spectra sequentially, eliminating the need to manually click the "Update" button for each entry. This streamlined workflow significantly enhances the efficiency of large-scale RNA/DNA MS/MS dataset analysis.

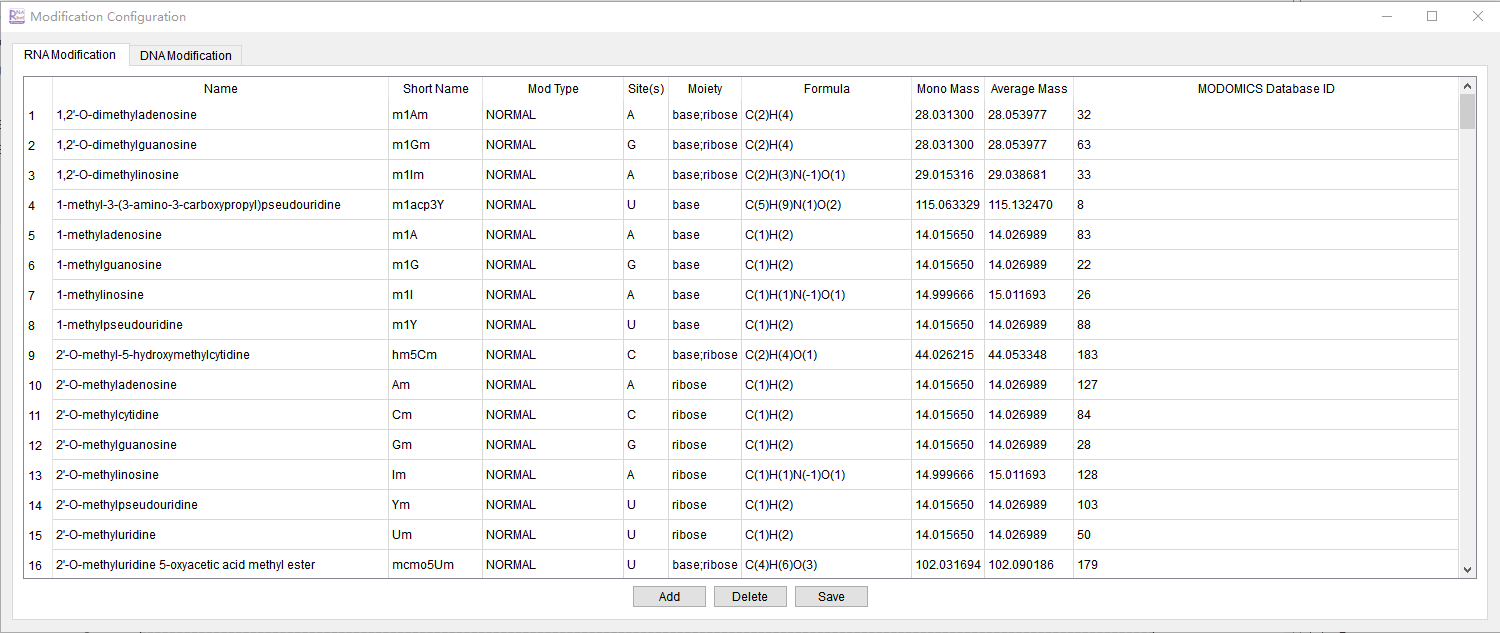

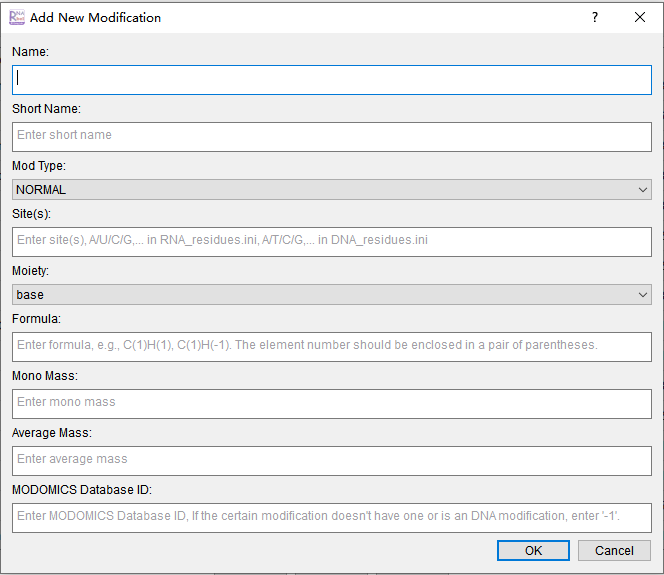

**A)**

**B)**

**Supplementary Figure S5: Customization GUI for RNA Modifications in RNabel**

This figure illustrates the graphical user interface (GUI) in RNabel for defining custom RNA modifications. To begin, users click "Modification Config" in the Tools menu, which opens the configuration interface shown in Panel A. In this interface, users can define custom RNA modifications by specifying the following parameters: **1) Name**: N6,2'-O-dimethyladenosine (We use **m6Am** as an example); **2) Short name**: m6Am; **3) Mod type**: NORMAL; **4) Sites**: A; **5) Moiety**: base;ribose; **6) Formula**: C(2)H(4); **7) Monoisotopic mass**: 28.0313; **8) Average isotopic mass**: 28.053977; **9) MODOMICS ID**: 82.

Once users click "Add", Panel B appears, prompting them to fill in all required fields for the new modification. The fields contain helpful tips for accurate data entry. After entering all information, users click "OK" in Panel B and "Save" in Panel A to save the new configuration. The successfully added modification will then be saved in the modification.ini file for future use. This interface allows users to define and store novel or uncommon RNA chemical modifications for precise spectrum annotation.

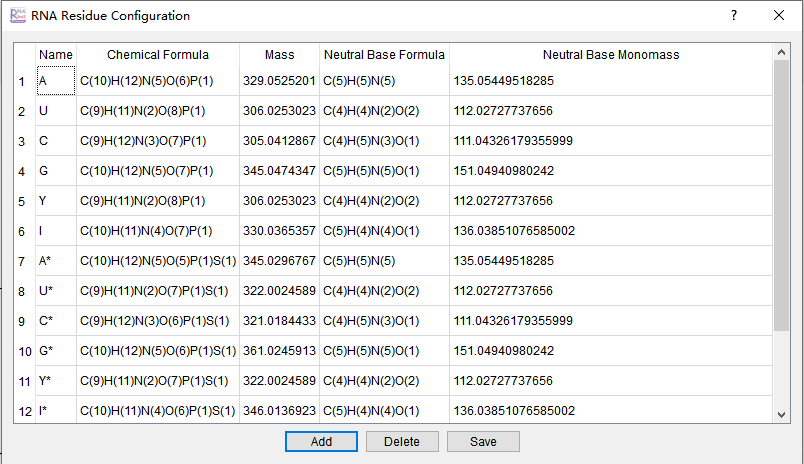

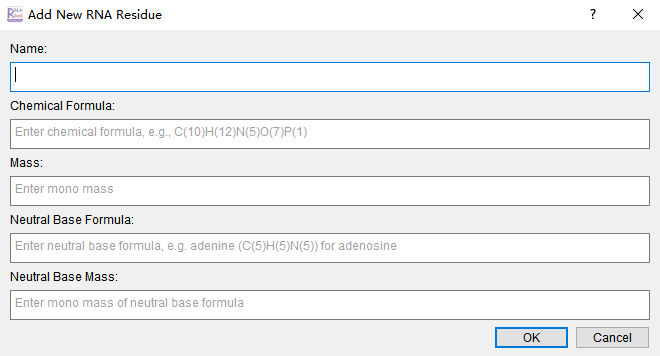

**A)**

**B)**

**Supplementary Figure S6: Customization GUI for Defining New Residue Types of RNA/DNA in RNabel**

This figure illustrates how users can define custom nucleotide residues through the RNabel GUI. Upon clicking the Residue Config option in the toolbar, users can view a window displaying all previously defined residues (subfigure A). To define a new residue, users click the Add button, which opens subfigure B. In subfigure B, users are prompted to enter the residue's name, chemical formula, monoisotopic mass, neutral base formula, and neutral base mass. (Here, the neutral base refers to the base in its uncharged form.) After inputting the required information, users click OK, and then save the new residue by clicking the Save button in subfigure A. The newly defined residue will appear in the list in subfigure A, confirming that the definition was successful. This process enables users to expand RNabel’s functionality by adding non-canonical nucleotide residues for their analysis.

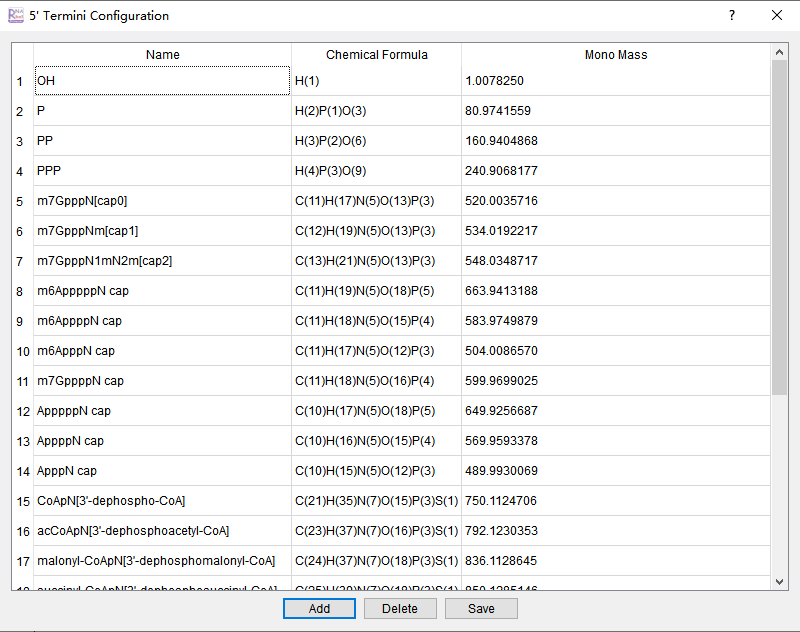

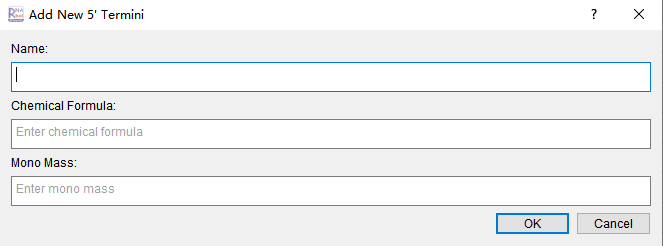

**A)**

**C)**

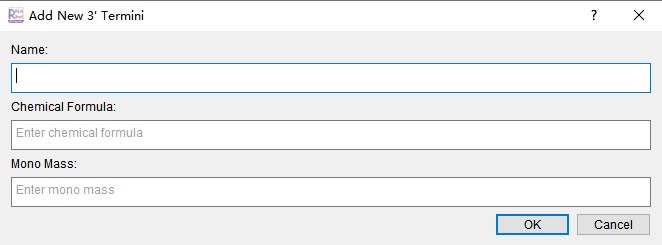

**B)**

**D)**

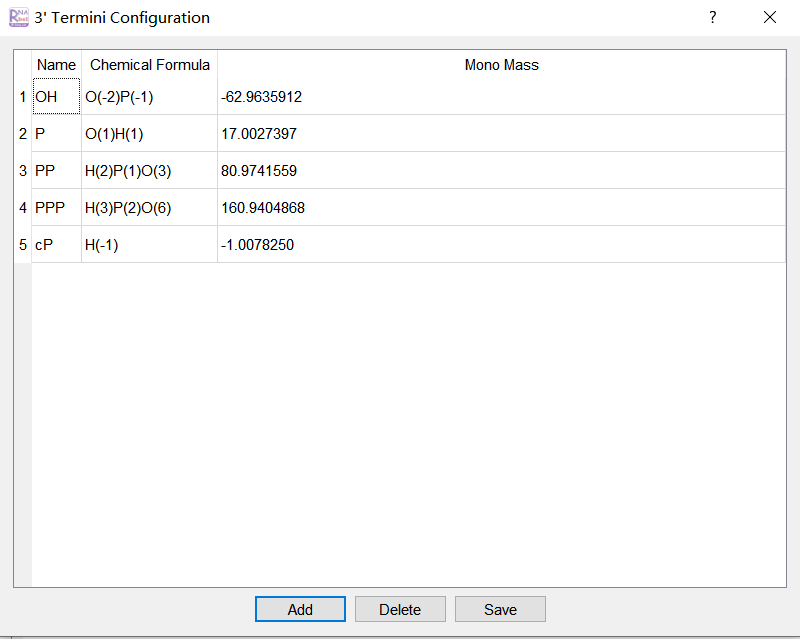

**Supplementary Figure S7: Customization GUI for Defining New Terminal Groups in RNabel**

This figure illustrates how users can define custom 5′ and 3′ terminal groups in RNabel. Upon selecting the Terminal Configuration option, users can view the list of previously defined 5′ terminal groups (subfigure A) and 3′ terminal groups (subfigure C). To define a new 5′ terminal group, users click the Add button, which opens subfigure B. In subfigure B, users are prompted to enter the name, chemical formula, and monoisotopic mass of the 5′ terminal group. After entering the required information, users click OK to confirm, and the newly defined group will appear in the list (subfigure A), indicating successful addition. The process for defining a 3′ terminal group is similar to that for the 5′ terminal group and is therefore not elaborated here. This customization feature allows users to incorporate novel terminal modifications that affect RNA/DNA sequence ends, thus enhancing the flexibility of RNA/DNA construct analysis and ensuring RNabel’s adaptability to various experimental setups.

**Supplementary Figure S8: Customization GUI for Defining New Sub-nucleotide fragments in RNabel**

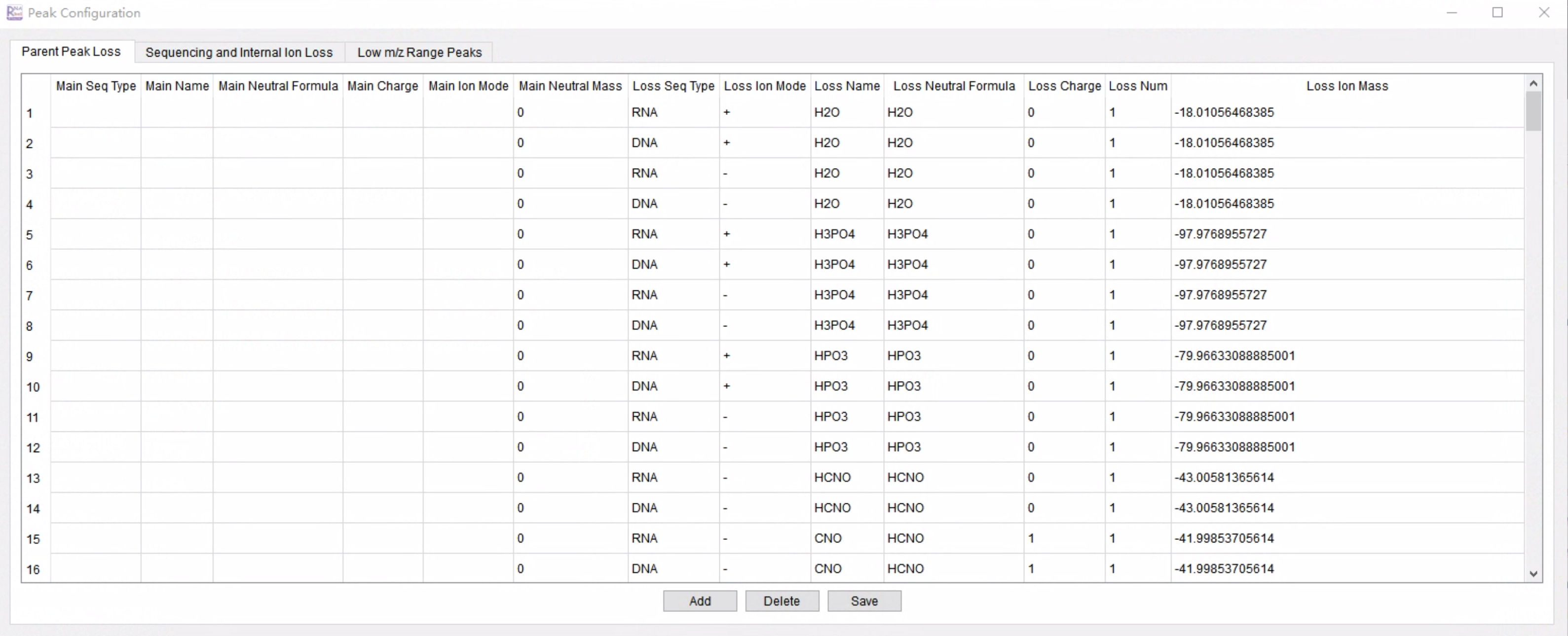

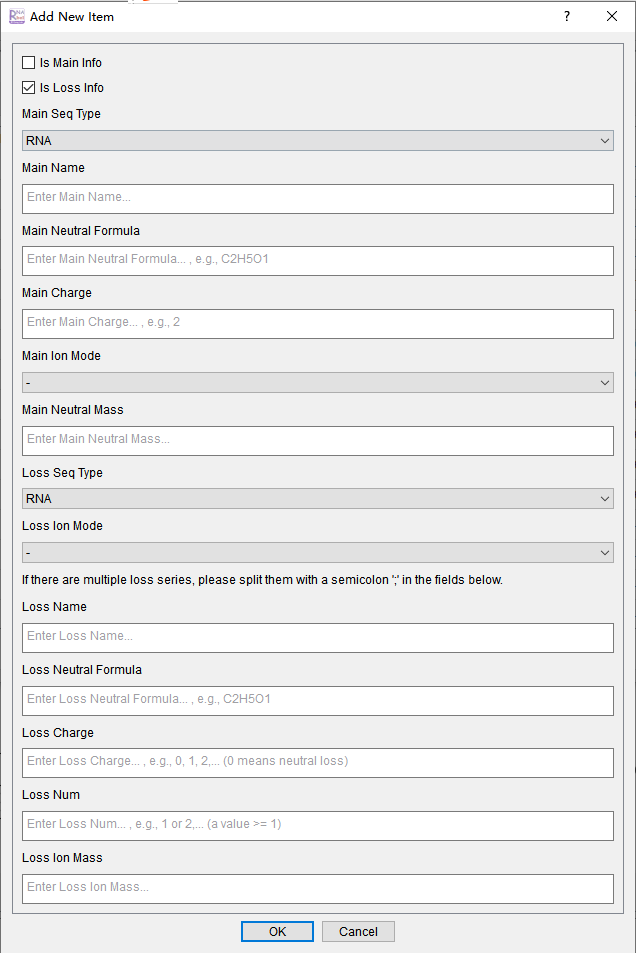

**A)**

**B)**

This figure illustrates how users can define custom sub-nucleotide fragments in RNabel. (A) The interface that appears when the Sub-nucleotide Fragment Config option is selected from the toolbar. In this window, users can view all previously defined sub-nucleotide fragments, which are the types of losses considered during spectrum annotation in RNabel. Sub-nucleotide fragment in RNabel are categorized into three types: 1) Parent Sub-nucleotide Fragment, *e.g.,* loss of a base or phosphate off a parent ion; 2) Sequencing and Internal Ion Loss, *i.e.,* loss of a small chemical group off a sequencing ion or internal ion; and 3) Low m/z Range Peaks, *i.e.,* fragment ions in the low m/z range. These losses are displayed in different tabs, with the current tab displaying the configuration interface for Sequencing and Internal Ion Loss. To define a new sub-nucleotide fragment, users can click the Add button, which opens the interface shown in (B).

In the window of Add New Item (B), users are prompted to input several parameters for the sub-nucleotide fragment definition. Each sub-nucleotide fragment is characterized by two components: Main Information and Loss Information. Parent ion losses and sequencing & internal ion losses should define the loss in the Loss Information section, with the sequence type and ion mode specified in the Main Information. Low m/z range peaks, though derived from other fragments, are treated as independent peaks and must be defined in the Main Information section. Both components require the sequence type (DNA/RNA), name, neutral formula, charge, and ion mode (positive or negative). The Loss Information section also includes additional parameters, such as loss number (loss num), allowing users to specify multiple simultaneous losses (e.g., two A bases and one U base lost from the same peak). For example, to define a sub-nucleotide fragment with two A bases and one U base, users input loss name = A; U, loss charge = 0; 0, and loss num = 2; 1, with neutral formula and ion mass automatically calculated. After inputting all necessary information, users click OK in the “Add New Item” window (B). Then, upon returning to the “Sub-nucleotide fragment Config” window (A), the new loss will appear in the list of defined sub-nucleotide fragments once the Save button is clicked.

It is important to note that the three categories of sub-nucleotide fragments (Parent loss, Sequencing & Internal ion loss, and Low m/z range peaks) are mutually exclusive and each has its own dedicated definition section. Specifically, for Low m/z range peaks, the Main Information should be used, as these peaks are derived from other fragments that have lost specific groups. In the “Add New Item” window (B), users should ensure that the Main Info checkbox is selected when defining these low m/z range peaks.

This customizable sub-nucleotide fragment feature allows for the annotation of complex fragmentation patterns and facilitates the exploration of novel chemical fragmentation events in RNA MS/MS spectra.

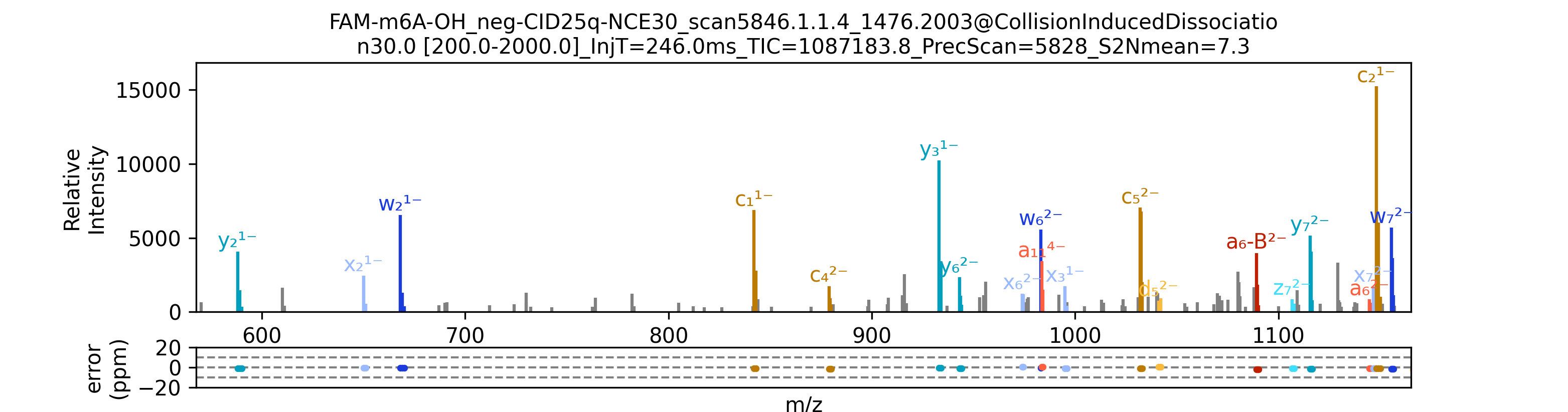

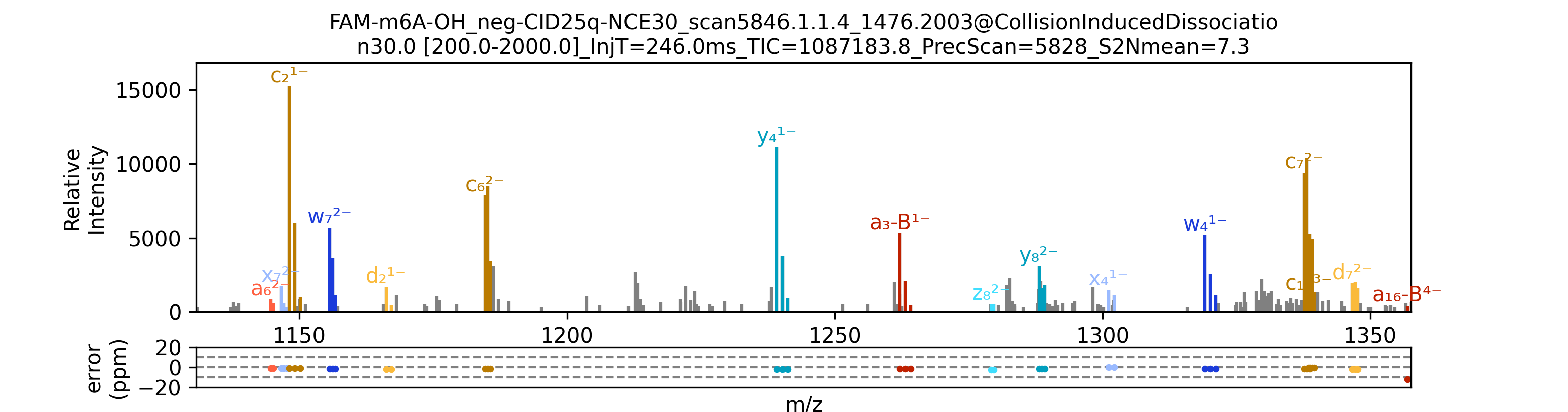

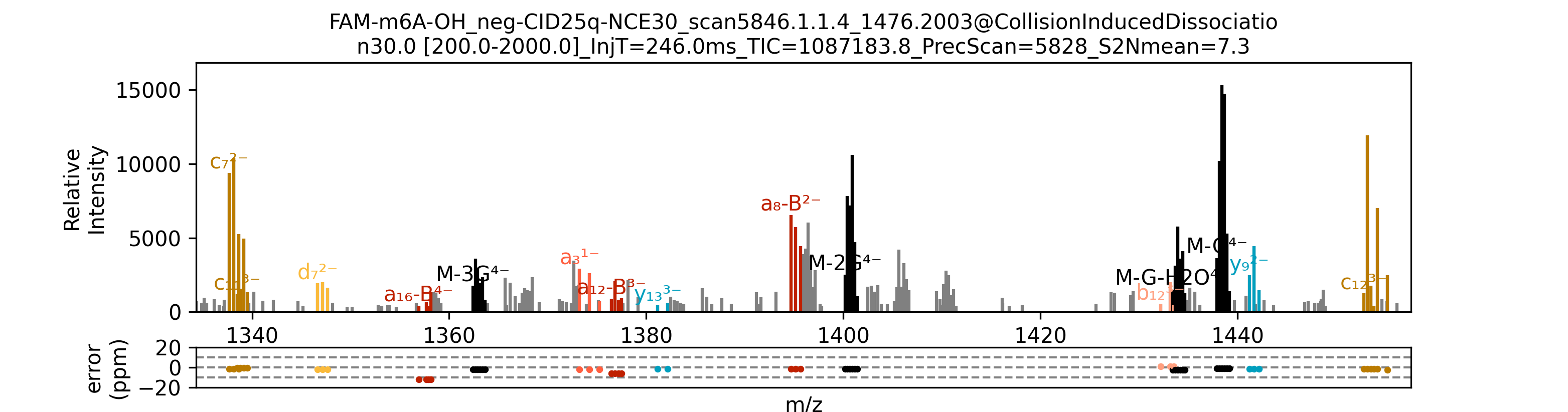

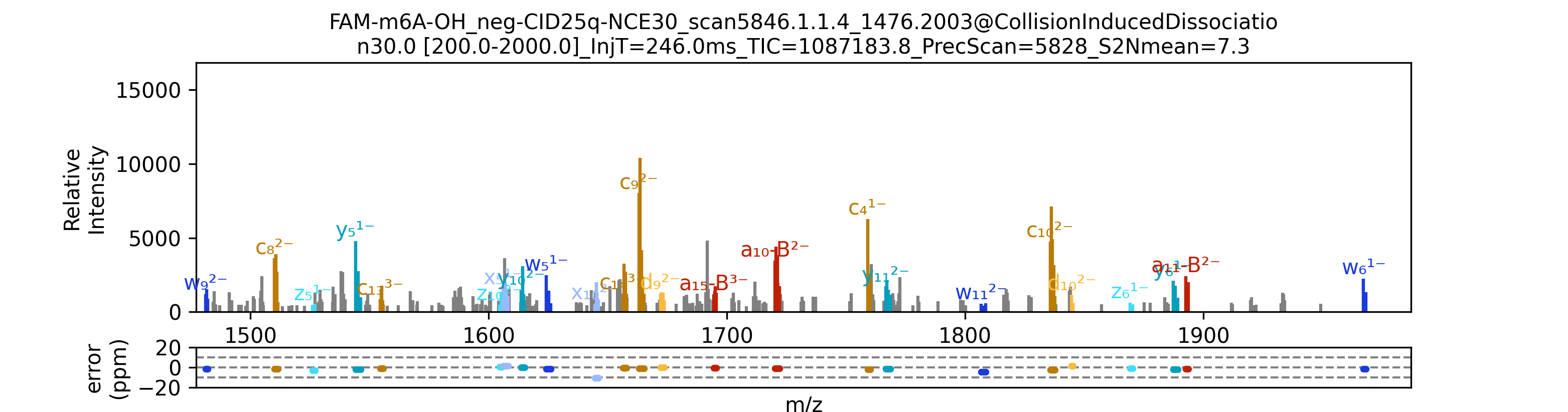

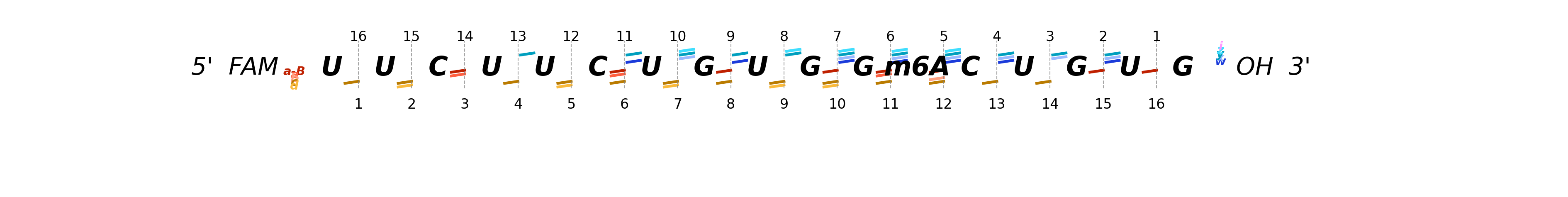

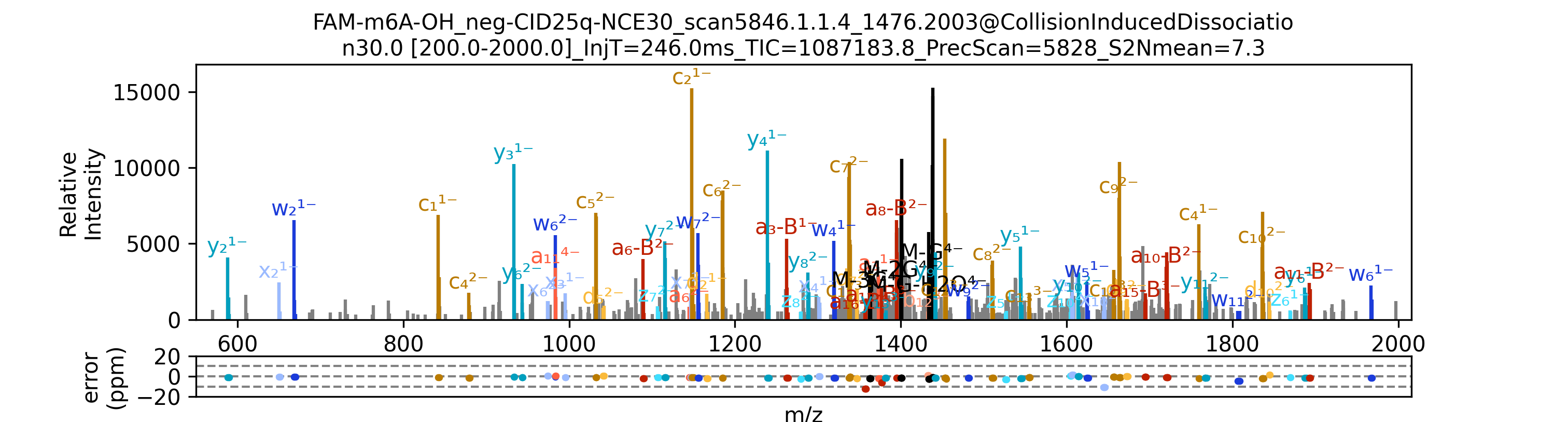

**Supplementary Figure S9: Detailed segment-by-segment zoomed-in views of the annotated MS/MS spectrum for the m⁶A-modified RNA oligonucleotide (FAM-m6A-OH, 5ʹ-FAM-UUCUUCUGUGG[m⁶A]CUGUG-OH-3ʹ), corresponding to the spectrum shown in Figure 6A.**

The top panel displays the full-range MS/MS spectrum, while the subsequent panels present consecutive zoomed-in *m/z* segments covering the entire spectral range, enabling detailed inspection of individual fragment ion assignments. Error plot for each annotated peak is shown in ppm below each segment. Data were acquired using collision-induced dissociation (CID) at a normalized collision energy (NCE) of 30.

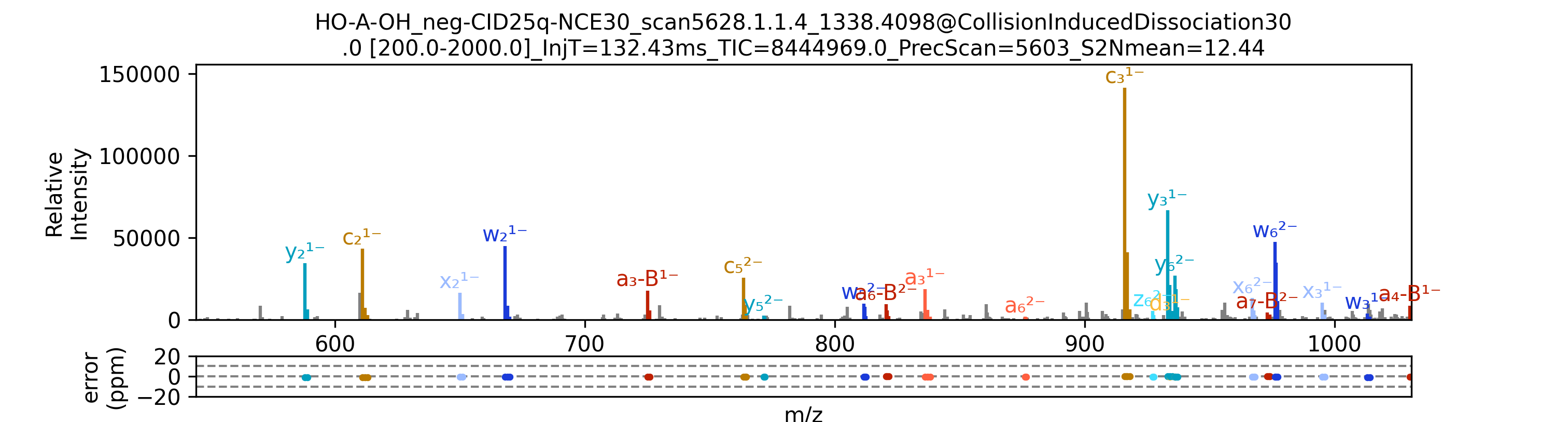

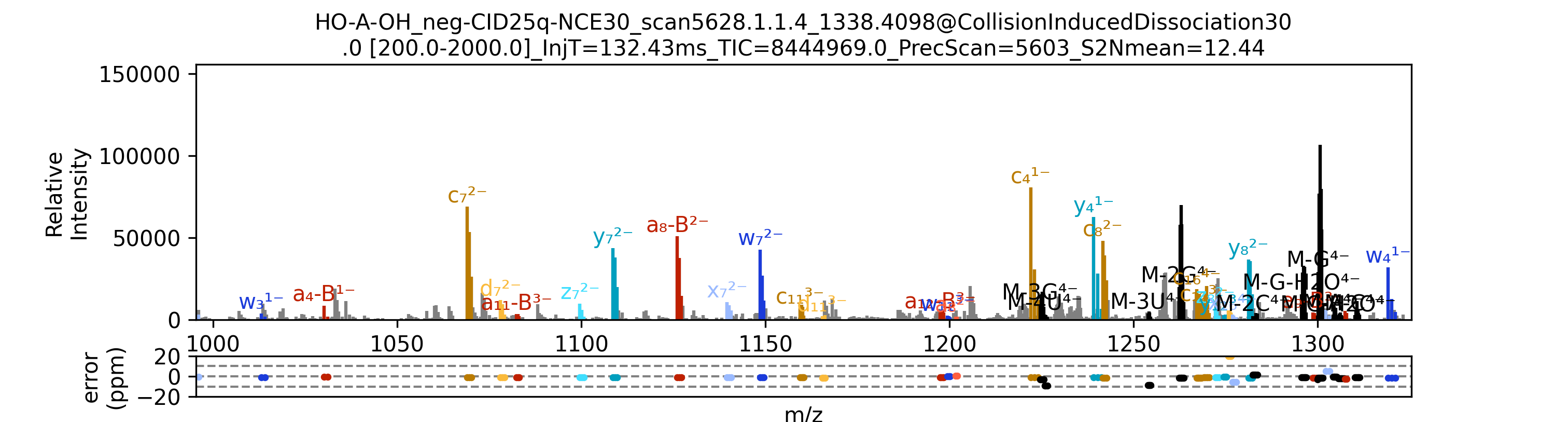

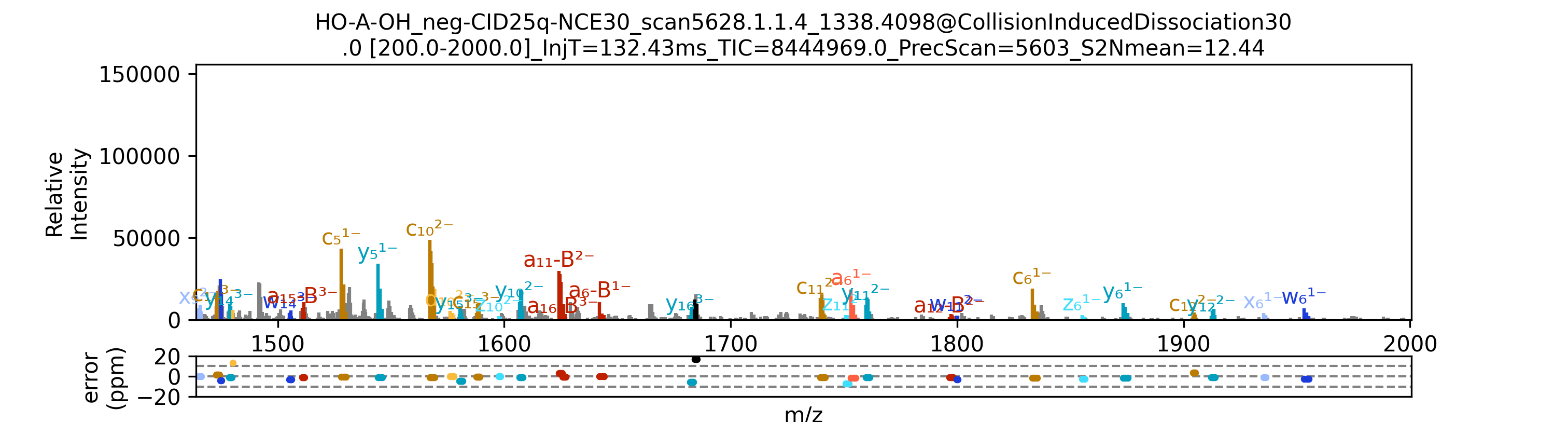

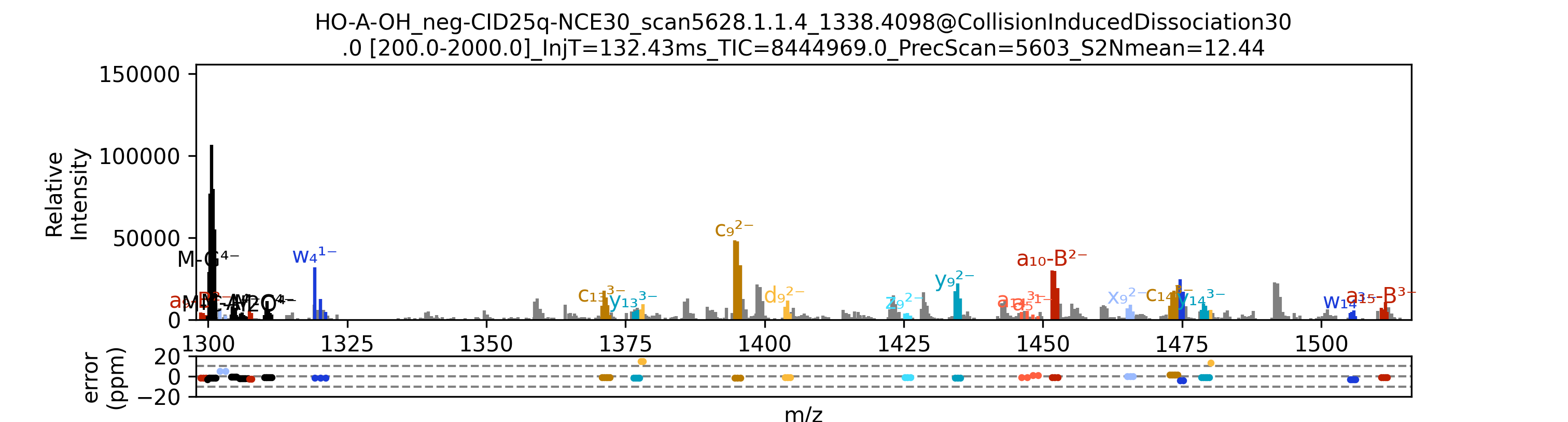

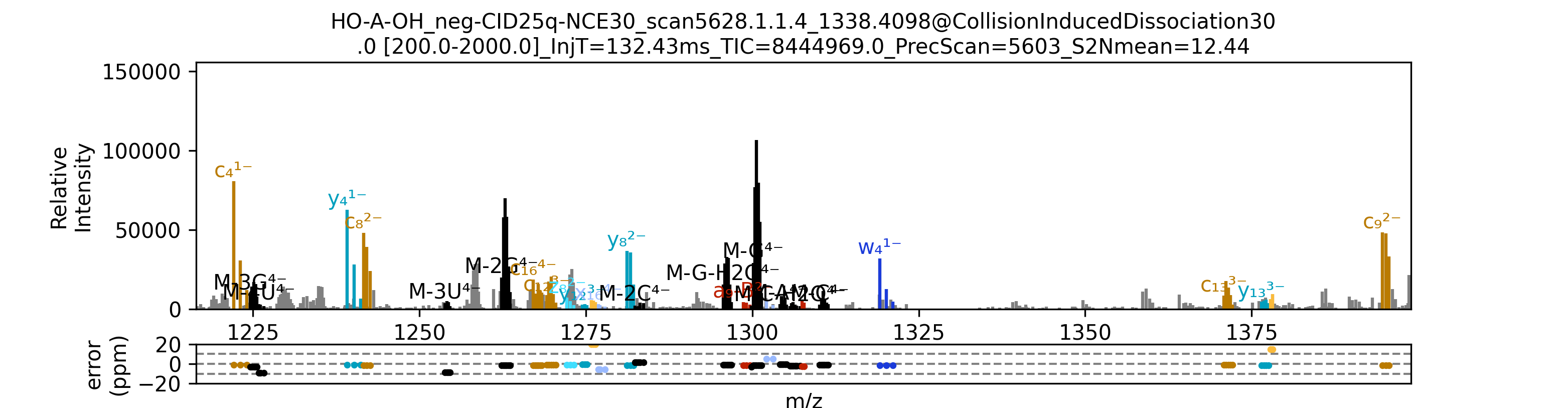

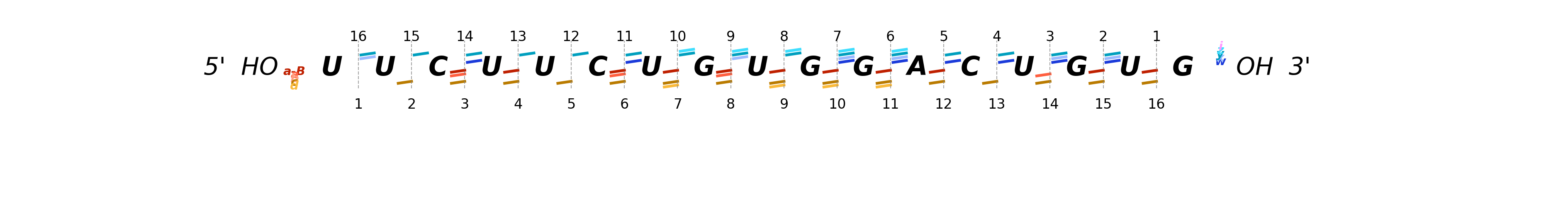

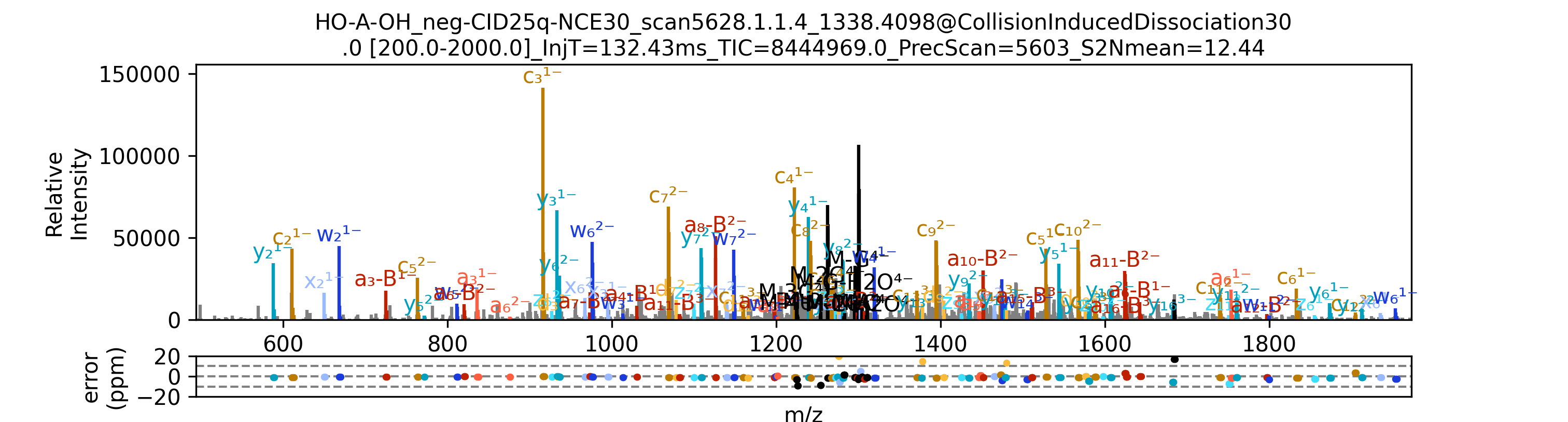

**Supplementary Figure S10: Detailed segment-by-segment zoomed-in views of the annotated MS/MS spectrum for the unmodified RNA oligonucleotide (HO-A-OH, 5ʹ-HO-UUCUUCUGUGGACUGUG-OH-3ʹ), corresponding to the spectrum shown in Figure 6B.**

The top panel displays the full-range MS/MS spectrum, while the subsequent panels present consecutive zoomed-in *m/z* segments covering the entire spectral range, enabling detailed inspection of individual fragment ion assignments. Data were acquired using collision-induced dissociation (CID) at a normalized collision energy (NCE) of 30.

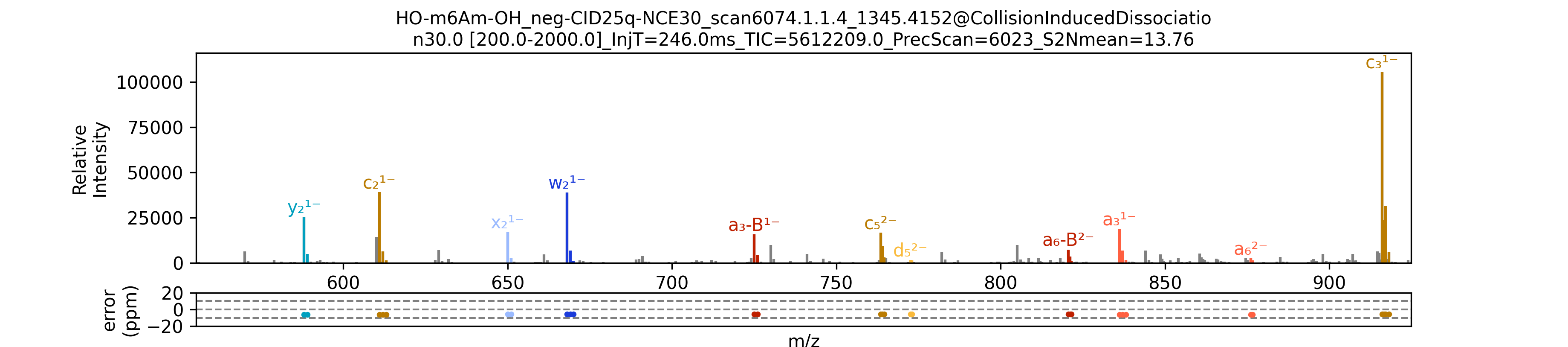

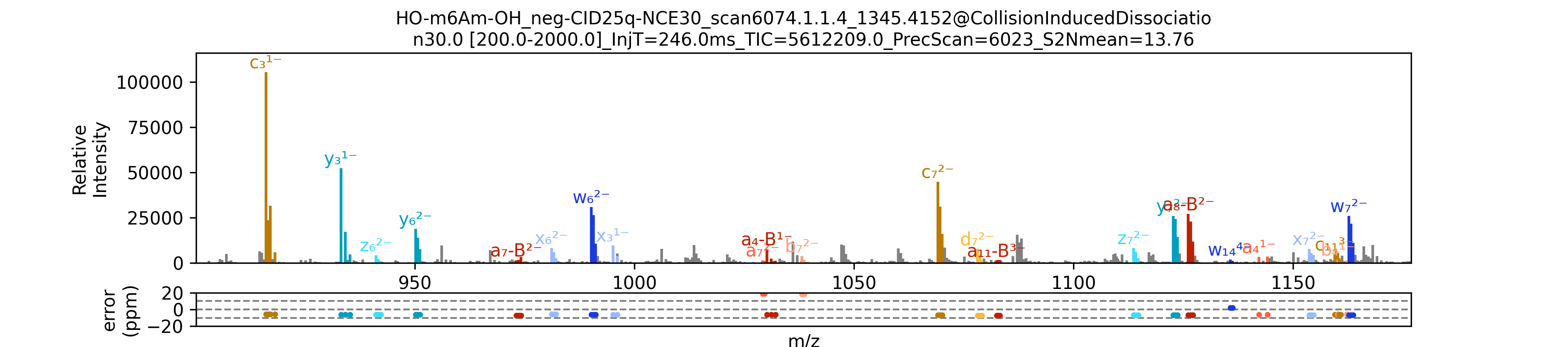

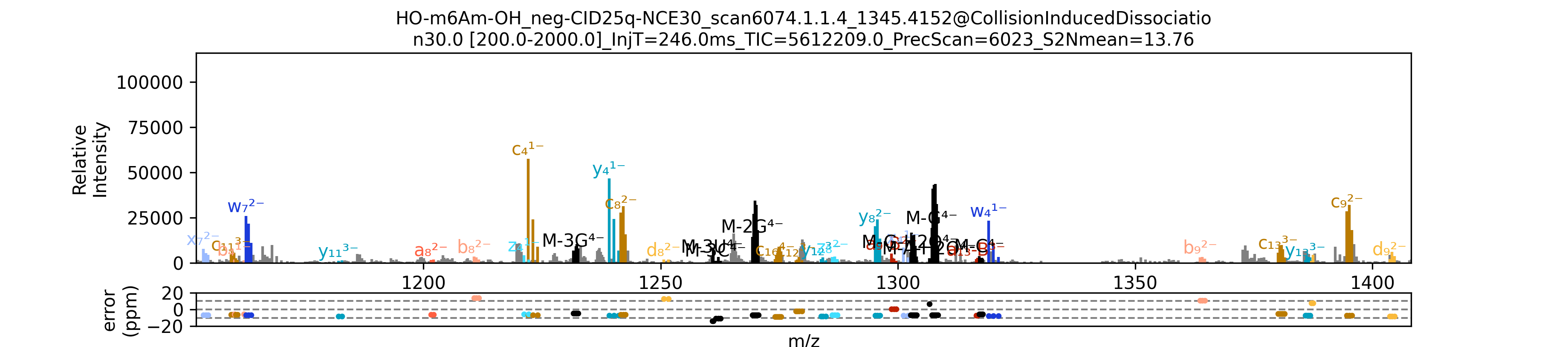

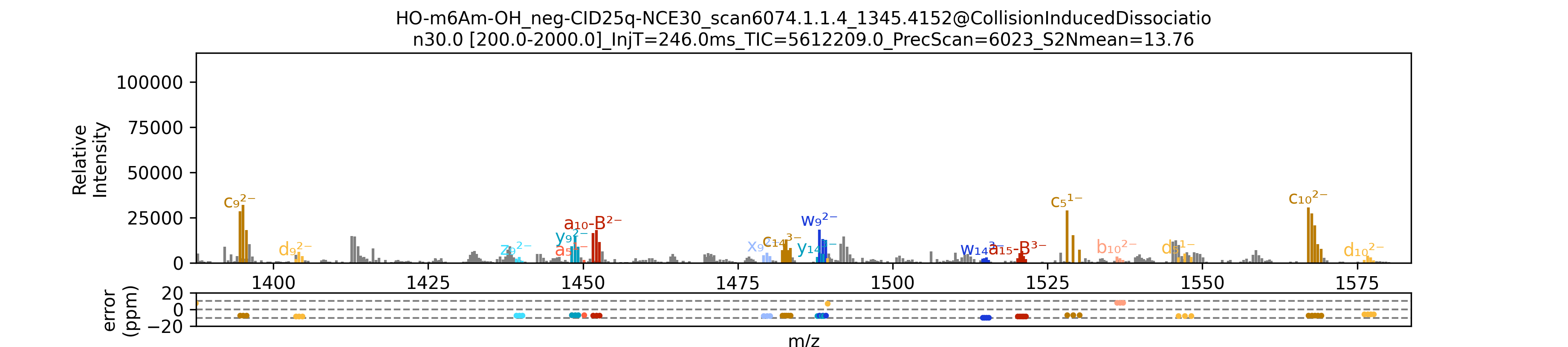

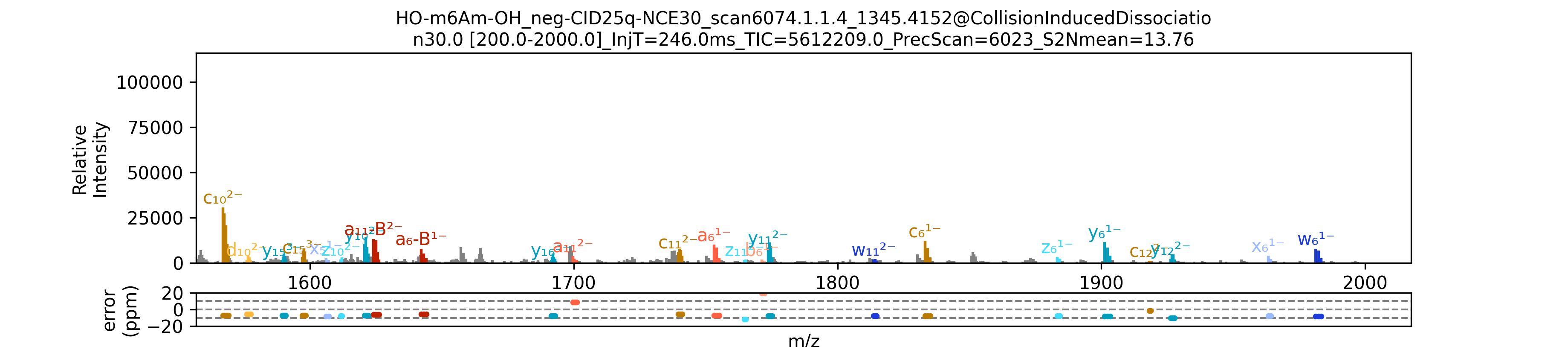

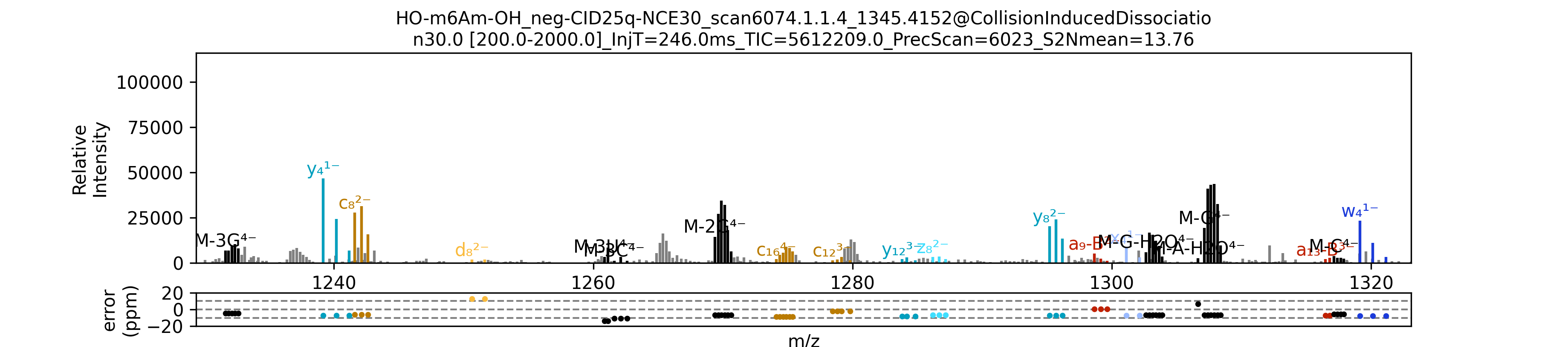

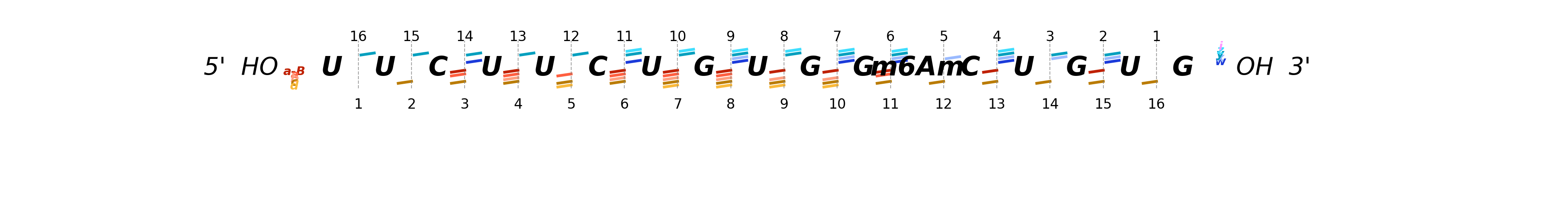

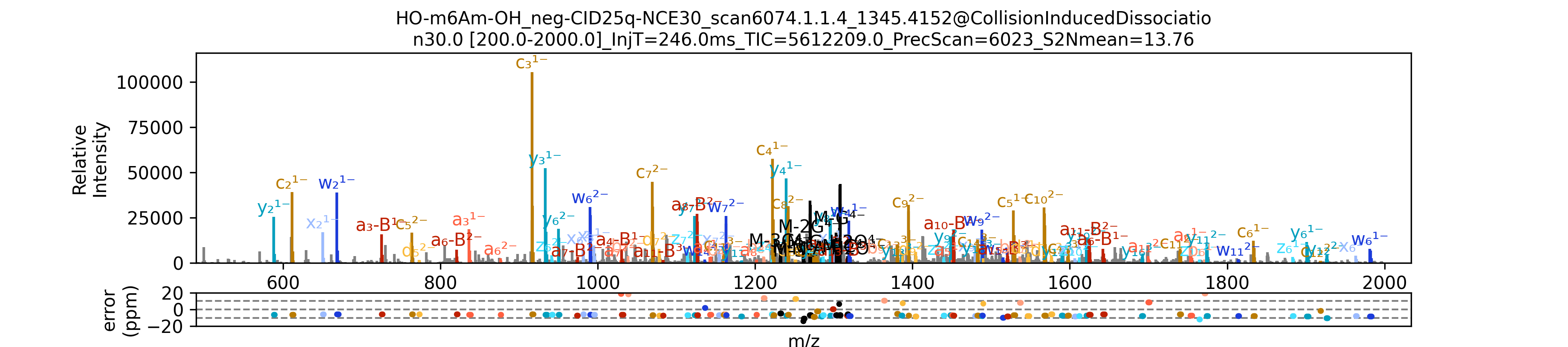

**Supplementary Figure S11: Detailed segment-by-segment zoomed-in views of the annotated MS/MS spectrum for the m⁶Am-modified RNA oligonucleotide (HO-m6Am-OH, 5ʹ-HO-UUCUUCUGUGG[m⁶Am]CUGUG-OH-3ʹ), corresponding to the spectrum shown in Figure 6C.**

The top panel displays the full-range MS/MS spectrum, while the subsequent panels present consecutive zoomed-in *m/z* segments covering the entire spectral range, enabling detailed inspection of individual fragment ion assignments. Data were acquired using collision-induced dissociation (CID) at a normalized collision energy (NCE) of 30.

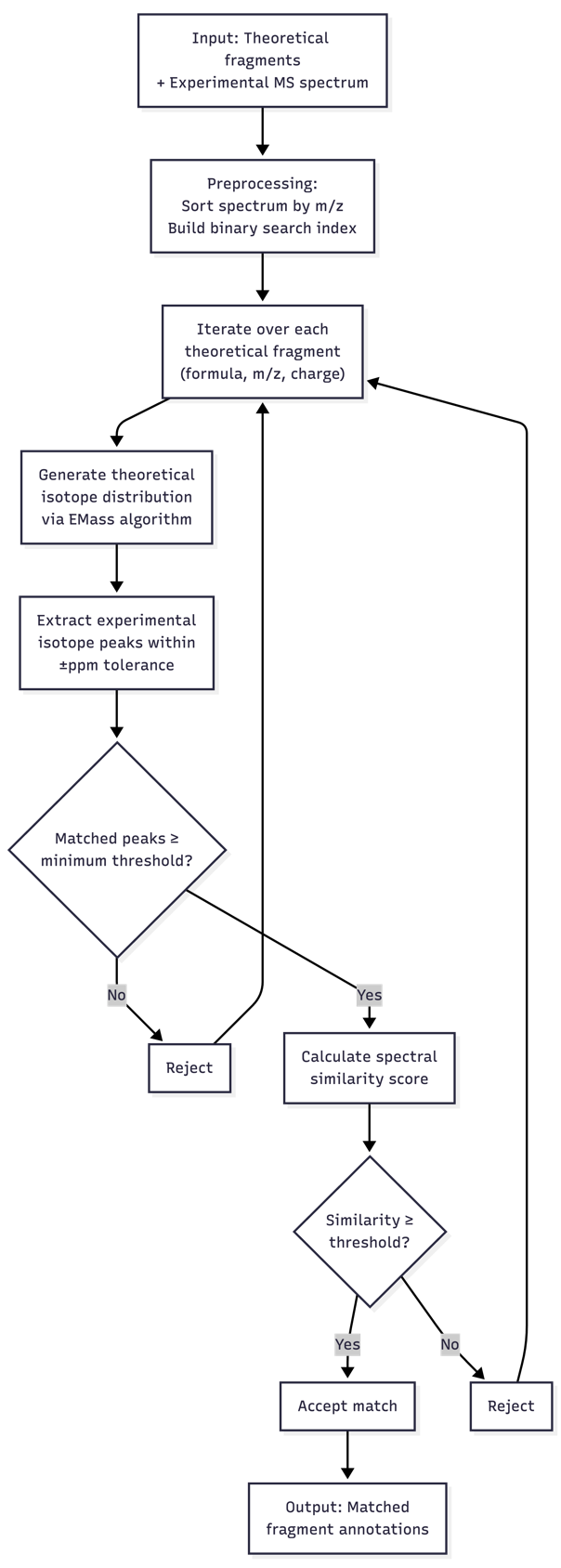

**Supplementary Figure S12. Workflow of the isotope distribution matching (IDM) algorithm.**

Theoretical isotopic distributions are generated for each candidate fragment, assuming a charge state from 1 to that of the precursor ion. A theoretical isotopic distribution is compared against the experimental one using cosine similarity scoring. Fragments passing both the minimum peak count and the similarity threshold are accepted as valid matches.

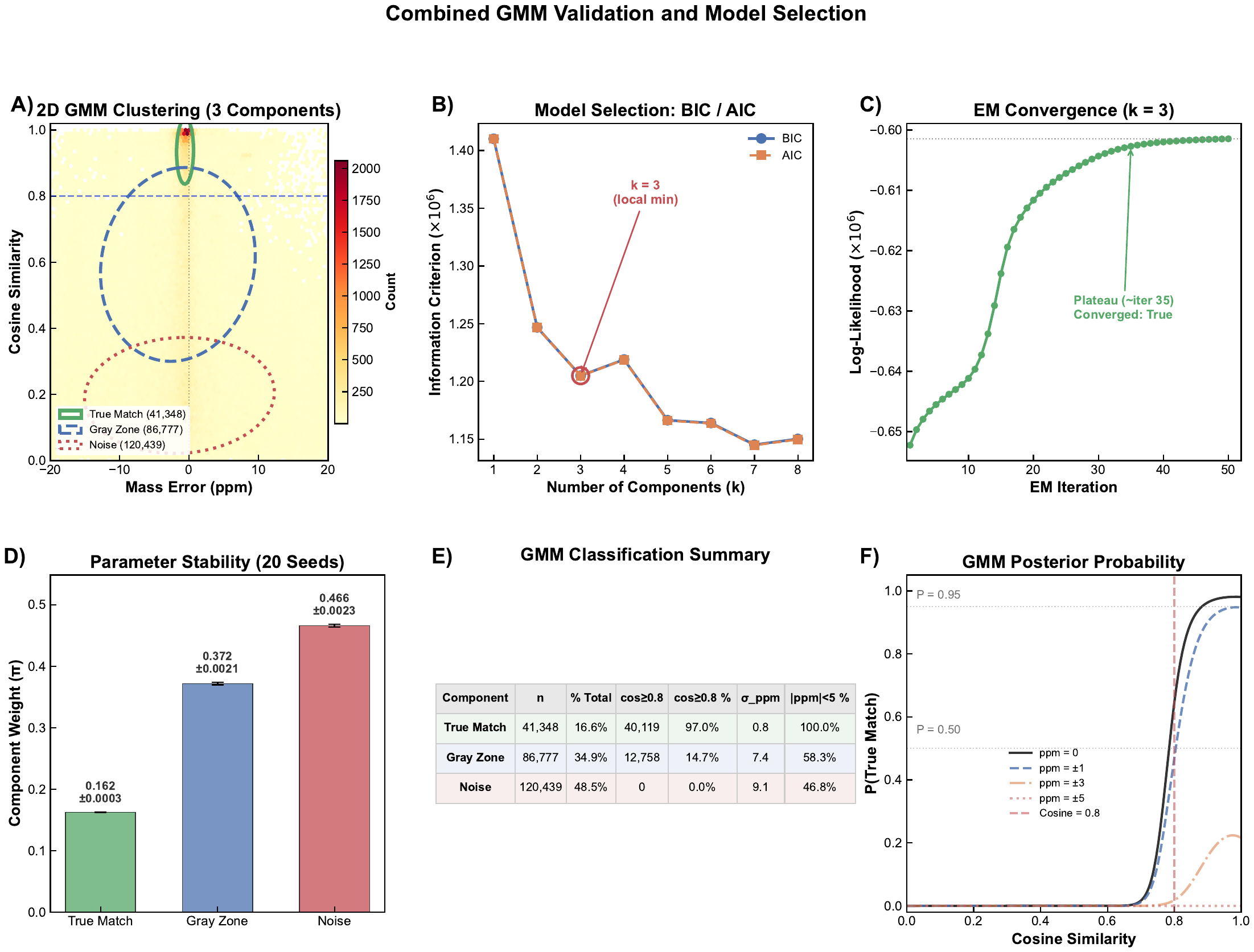

**Supplementary Figure S13. Unsupervised Gaussian Mixture Model (GMM) for determining the cosine similarity threshold.**

(A) Two-dimensional density map of match candidates (n = 248,564; |ppm| ≤ 20) in the mass error versus cosine similarity space, overlaid with 1.5σ covariance ellipses from a three-component GMM fitted to the joint feature space of cosine similarity and signed ppm error. The model autonomously identified three physically interpretable populations: True Match (n = 41,339; high cosine, narrow ppm), Gray Zone (n = 86,932; moderate cosine, intermediate ppm), and Noise (n = 120,293; low cosine, diffuse ppm). The blue dashed line indicates cosine = 0.8, which coincides with the boundary between the True Match and Gray Zone ellipses. (B) Bayesian information criterion (BIC, blue circles) and Akaike information criterion (AIC, orange squares) as a function of the number of GMM components (k = 1–8). A local BIC minimum was observed at k = 3 (BIC = 1.205 × 10⁶), with BIC increasing upon addition of a fourth component (k = 4, BIC = 1.219 × 10⁶), indicating that three macro-populations constitute the natural grouping of the data. While higher k values (≥5) achieved progressively lower BIC by capturing intra-population substructure, the three-component model was selected based on parsimony and physical interpretability. (C) Expectation-maximization (EM) convergence trace for the k = 3 model, showing log-likelihood as a function of iteration number. The algorithm reached a stable plateau within ~35 iterations, confirming robust convergence. (D) Component weight stability across 20 independent random initializations (seeds 0–19). Error bars represent ±1 standard deviation. The negligible variability (σ < 0.004 for all components) demonstrates that the three-component solution is unique and independent of initialization. (E) GMM classification summary. Of the 41,339 True Match candidates identified by GMM, 97.0% exhibited cosine similarity ≥ 0.8, with a ppm standard deviation of 0.8 (consistent with instrument mass accuracy). None of the 120,293 Noise candidates exceeded the 0.8 threshold. Data filtered to |ppm| ≤ 20 (n = 248,564) (F) GMM posterior probability of belonging to the True Match component, P(True Match), as a function of cosine similarity at fixed mass errors of ppm = 0 (black solid), ±1 (blue dashed), ±3 (orange dash-dot), and ±5 (red dotted). At cosine = 0.8 with ppm = 0, P(True Match) ≈ 0.65; P(True Match) exceeds 0.95 at cosine ≈ 0.9. At ppm = ±5, P(True Match) remains below 0.2 even at cosine = 1.0, demonstrating that cosine similarity and mass accuracy function as orthogonal quality dimensions.

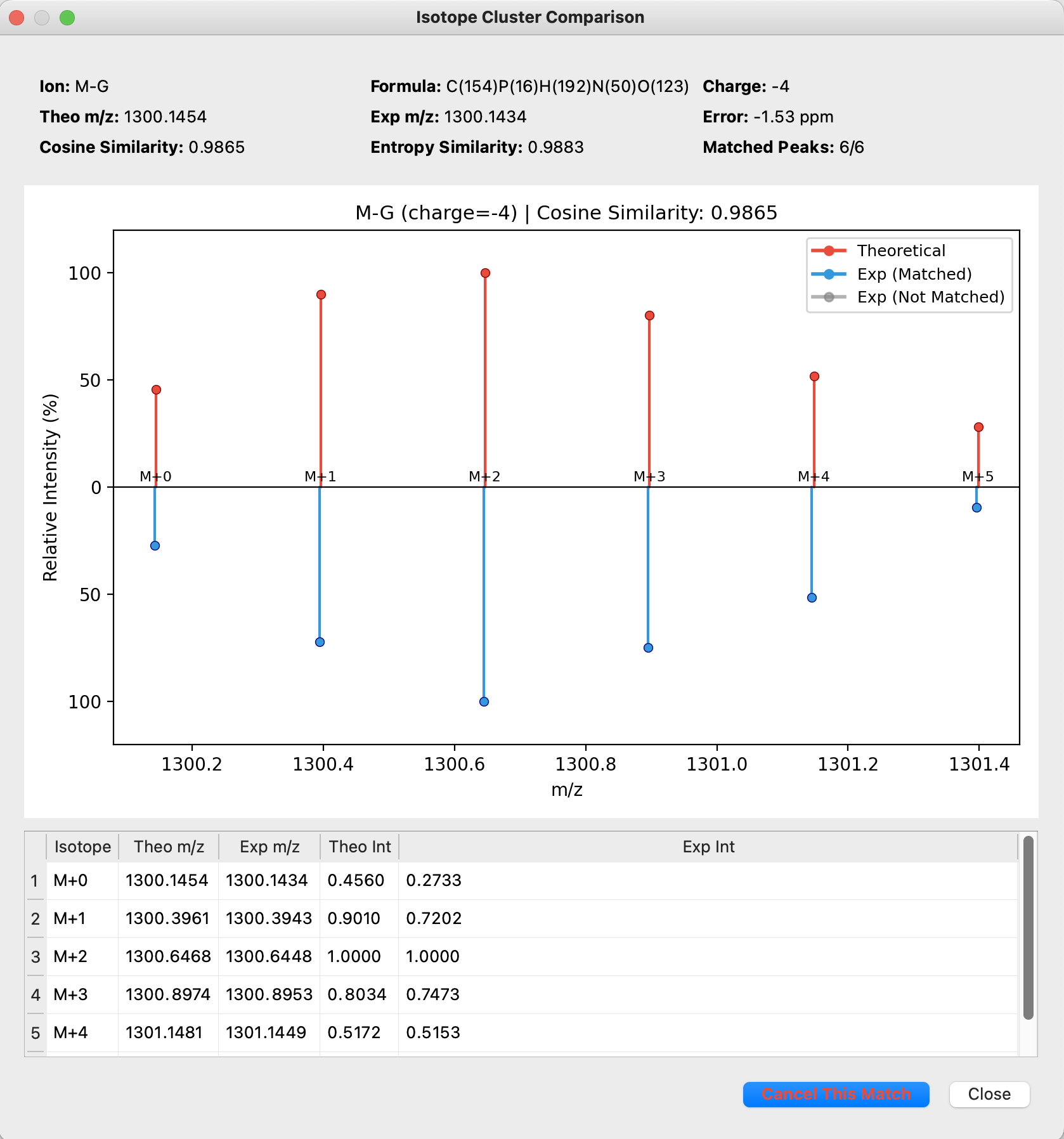

**Supplementary Figure 14. Interactive mirror-plot interface for inspecting a match between two isotopic distributions.**

Example showing the isotopic distribution match for a fragment ion (charge state: −4). The upper panel displays the theoretical isotopic distribution (red), and the lower panel shows a segment of the experimental spectrum in which matched peaks are highlighted in blue and unmatched peaks in gray. The table below lists the individual isotopic peaks with their theoretical and experimental m/z values and intensities. Cosine similarity and spectral entropy similarity scores are displayed to assist users to evaluate the quality of match. Users can accept or reject each match through this interface.

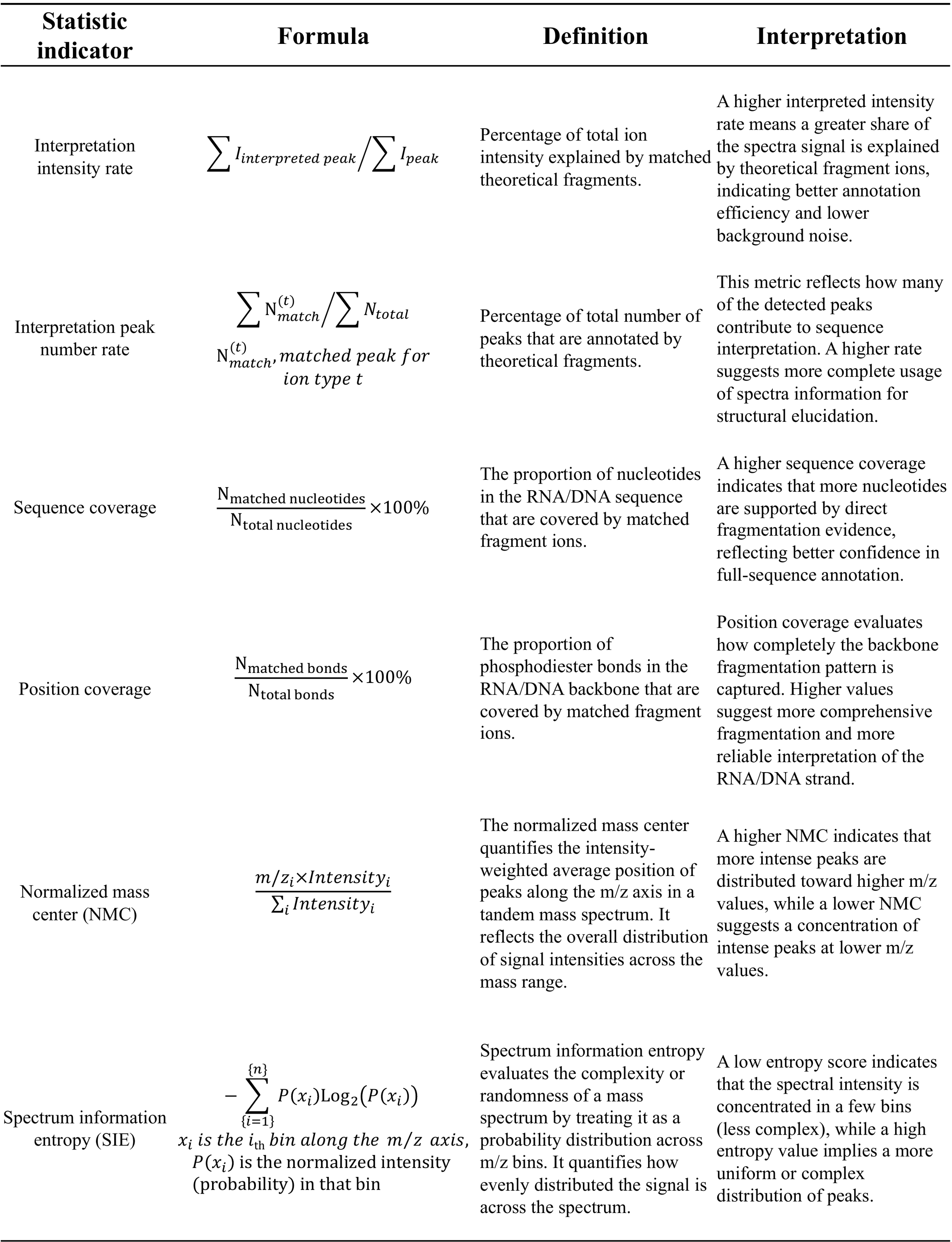

**Supplementary Table S1: Statistical metrics calculated by RNabel to evaluate tandem MS spectrum annotation quality**

This table summarizes the definitions of six statistical indicators calculated by RNabel. Sequence coverage is the proportion of nucleotides in the RNA sequence that are supported by matching fragment ions. Position coverage is the proportion of phosphodiester bonds in the RNA backbone that are supported by matching fragment ions. Interpreted intensity rate is the proportion of total spectral intensity explained by matched fragment ions. Interpreted peak rate is the proportion of total spectral peaks that are assigned to matched fragment ions. Normalized mass center (NMC) quantifies the intensity-weighted average position of peaks along the m/z axis, reflecting the overall distribution of spectral intensity. Spectrum information entropy (SIE) measures the complexity of the spectrum based on the probability distribution of peak intensities across m/z bins, indicating how evenly distributed the signal is.

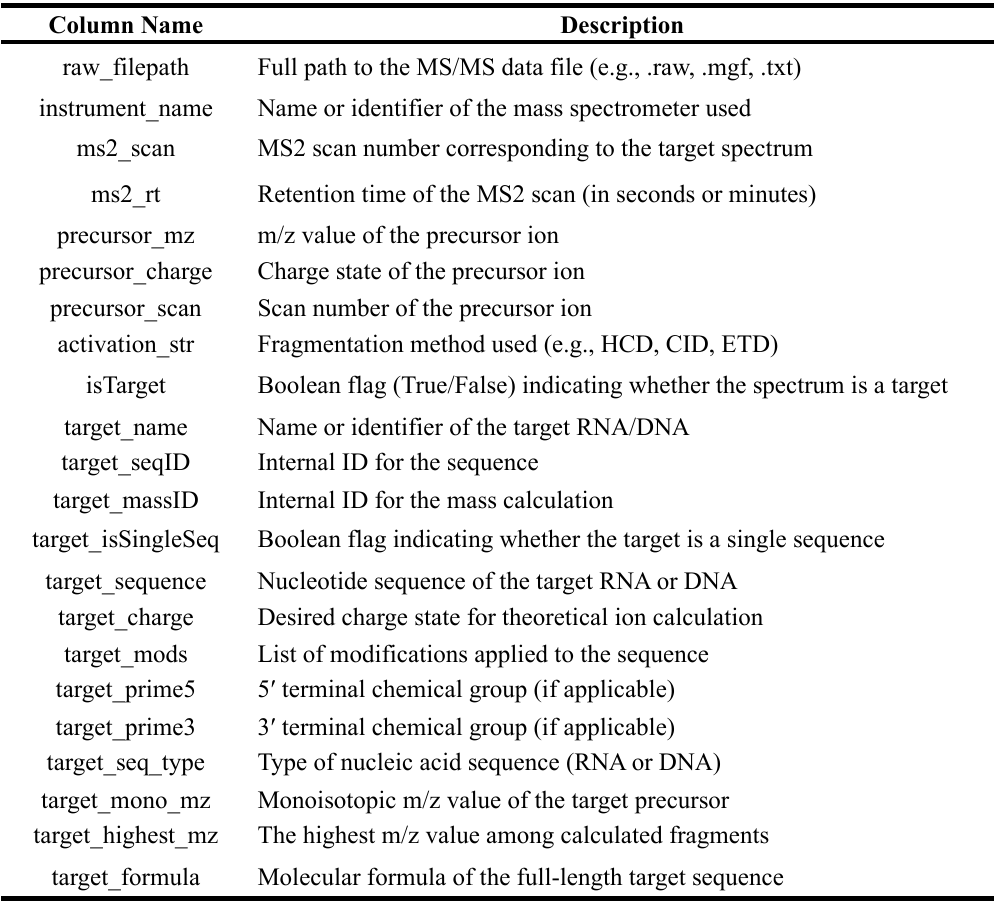

**Supplementary Table S2: Required columns in RNabel batch input files**

This table lists the required column headers and their descriptions for the RNabel batch input file format. Each row in the batch file corresponds to a single MS/MS spectrum and its associated sequence and metadata, enabling RNabel to efficiently perform automated multi-spectrum annotation.

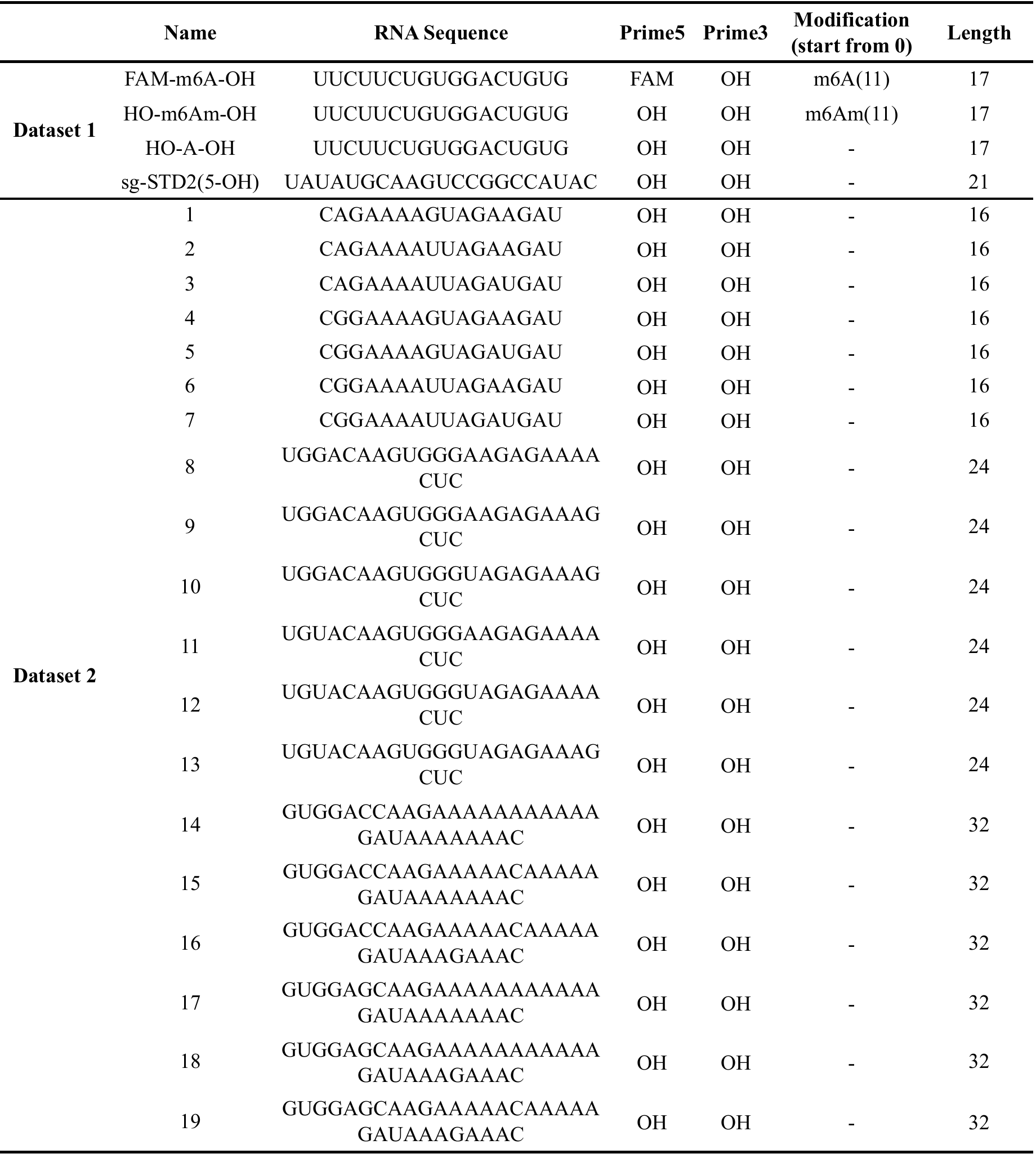

**Supplementary Table S3: Synthetic RNA oligos used in the development of RNabel.**
